# Sticky salts: overbinding of monovalent cations to phosphorylations in all-atom forcefields

**DOI:** 10.1101/2025.08.28.672842

**Authors:** Jules Marien, Julie Puyo-Fourtine, Chantal Prévost, Sophie Sacquin-Mora, Elise Duboué-Dijon

## Abstract

Phosphorylation is a major post-translational modification, which is involved in the regulation of the dynamics and function of Intrinsically Disordered Proteins (IDPs). We recently characterized a phenomenon, which we termed *n*-Phosphate collaborations (*n*P-collabs), where bulk cations form stable bridges between several phosphoresidues in all-atom molecular dynamic simulations. *n*P-collabs were found to be sensitive to the combination of forcefields and cation types. Here, we attempt to assess the physical relevance of these *n*P-collabs by evaluating the strength of the cation/phosphate interaction through osmotic coefficient (*ϕ*) calculations on the model 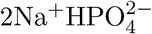 and 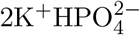 salts, using different classical forcefields for phosphorylations. All force-fields were found to overestimate the strength of the interaction to various degrees. We thus designed new parameters for CHARMM36m and AmberFF99SB-ILDN using the Electronic Continuum Correction (ECC) approach, which provide remarkable agreement for *ϕ* values for both cation types and over a range of concentrations. We provide a preliminary test of these ECC parameters for phosphorylations by simulating the sevenfold-phosphorylated rhodopsin peptide 7PP and comparing secondary chemical shifts to experimental data. Conformational ensembles resulting from the ECC-derived phosphorylated forcefields display both qualitative and quantitative improvements with regard to full-charge forcefields. We thus conclude that long-lasting *n*P-collabs are artifacts for classical forcefields born from the lack of explicit polarization, and propose a possible computational strategy for the extensive parameterization of phosphorylations. The presence of long-lived *n*P-collabs in simulations produced using classical forcefields is therefore a serious concern for the accurate modelling of multiphosphorylated peptides and IDPs, which are at the center of research questions regarding neurodegenerative diseases such as Alzheimer’s or Parkinson’s.

## INTRODUCTION

The last decades have seen a growing interest for intrinsically disordered proteins (IDPs) or intrinsically disordered regions (IDRs), ^1^ as they are central players for cellular function, notably for signalling, regulation and organization,^2–5^ and are also involved in numerous neurodegenerative diseases. ^6,7^ Unlike their folded counterparts, IDPs cannot be properly characterized by a single structure but should rather be described by a conformational ensemble, and this conformational variability is critical for their function.^8^ IDPs conformational ensembles and intermolecular interactions, and therefore their function, can be regulated by post-translational modifications (PTMs).^9–14^ Amongst them, phosphorylations are the most common PTMs^15^ and are known to preferentially target IDPs and IDRs.^16,17^

In addition to experimental techniques such as small-angle X-ray scattering (SAXs), nuclear magnetic resonance (NMR), circular dichroism or Förster resonance energy transfer (FRET),^18–22^ one can also use computer simulations to explore and characterize the conformational space of IDPs, and numerous force fields have been specifically parameterized to model disordered systems.^23–27^ However, the impact of phosphorylations on an IDP conformation and dynamics has been found previously in all-atom molecular dynamics simulations to strongly depend on the nature of the employed counterion (K^+^ or Na^+^) and on the chosen force field.^28,29^ In particular, different force fields yield drastically different pictures of the amount of ion pairing to phosphorylation and formation of cation mediated bridges (termed *n*P-collabs), which then reflect on the IDP physical properties, such as its radius of gyration and flexibility.^29–32^ Each phosphorylated moiety bears a −2 negative charge at pH 7. ^33^ The formation of ion pairs, especially with multiply charged species, is known to be particularly challenging for molecular simulations that are prone to overbinding artefacts. ^34–38^ These artefacts impact cation binding to protein, nucleic acids and phospholipids but also salt bridge formation in all non-polarizable force fields.

Here, we thus assess the physical relevance of these contrasting ion pairing behavior in phosphorylated IDPs using a minimal model of a phosphorylation, the monoprotonated phosphate 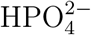, for which experimental data pertaining to ion pairing is available. Namely, we compare experimental osmotic coefficients of K_2_HPO4 and Na_2_HPO4 solutions with those resulting from calculations using six combinations of protein, ion and water force fields that we found to be commonly used in the IDP simulation literature. This comparison reveals a systematic overbinding of monovalent ions to phosphorylation.

Several avenues are being actively explored in the literature to overcome such artefacts, that are known to originate from the lack of electronic polarization in the most common biomolecular force fields. ^37^ One approach is polarizable force fields, that explicitly add extra degrees of freedom to account for electronic polarization.^39–41^ Development and validation of polarizable force fields for phosphorylated proteins is a very active field of research.^42,43^ Yet, polarizable force fields are still computationally more expensive and more complex to parameterize and use than standard ones, which led the community to explore alternative implicit strategies, where the effects of electronic polarization are included in a mean-field approach, within the standard functional terms of non-polarizable force fields. Several approaches can be found in the literature, including the parameterization of pair-specific Lennard-Jones parameters (often called the NBFIX approach using the CHARMM vocabulary) ^44^ or the derivation of implicitly polarized charges.^45–47^

In this work, we specifically explore the Electronic Continuum Correction (ECC) strategy, originally suggested by Leontyev and Stuchebrukhov ^46^ and increasingly popular in recent years for a variety of systems. The ECC is a physics-based approach that incorporates electronic polarizability as a dielectric continuum, which is equivalent to scaling the charge of charged moieties by a factor 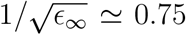, where *ϵ*_*∞*_ is the fast electronic component of the dielectric constant.^46,48^ This transferable approach has been shown, with contributions of two of us, to drastically improve the description of ion pairing in molecular simulations, ^37^ both in simple electrolytes,^49–51^ including with phosphates,^52,53^ and in biomolecular simulations of proteins, including, very recently, ion channels, ^54–56^ membranes,^36^ nucleic acids,^56,57^ and sulfated polysaccharides.^58^ Here, we investigate whether an ECC correction for the −2 charged phosphorylated groups can improve ion pairing of phosphorylated IDPs, and then characterize the impact of this modification on the physical properties of phosphorylated IDPs.

## MATERIAL AND METHODS

### Forcefields

Four main combinations of forcefields were employed in this study, that were previously used in the literature to study phosphorylated IDPs. We name A18 the association of AmberFB18^59^ with the TIP3P-FB water model^60^ and ionic parameters SPC/E from Joung and Cheatham (called later JC ions);^61^ A19 the association of AmberFF19SB^62^ with the OPC water model^63^ and ions from Sengupta et al. ^64^; A99 the association of AmberFF99SB-ILDN^65^ with TIP4P-D^66^ and ion parameters derived by Beglov and Roux ^67^ (called later BR ions); and C36 the association of CHARMM36m with its associated TIP3P and BR ions.^68^ Additional ionic parameters from the DES-Amber SF1.0 forcefield ^56^ were also tested in combination with A99. Ionic parameters are summarized in Table SI-1. For parameters of the phosphorylations, C36 contains them already; A18 is technically the name of the phosphorylation extension of forcefield AmberFB15;^69^ A19 was equiped with phosaa19SB; ^70^ and A99 with phosaa10^71^ with improvements from Steinbrecher et al..^72^

Common non-polarizable force fields, such as those described above, are known to suffer from artefactual overbinding for ion pairing due to the lack of electronic polarizability.^37^ Other forcefield families such as GROMOS simply transferred the charges of ATP nucleotide to phosphoresidues without further investigation of phosphate/cation interactions, ^73^ while the OPLS family saw a recent update by Usher et al.^14^ of the continuum solvent model ABSINTH. Their improved parameters notably had to be restricted to monoprotonated phosphoresidues since deprotonated forms were subject to *n*P-collabs deemed too stable by the authors (see Figure S1 of Ref^14^). Here, we test the so-called Electronic Continuum Correction (ECC) strategy to improve the ion pairing behavior of phosphorylated IDPs. ECC is a mean-field theory that implicitly takes into account electronic polarizability through a dielectric continuum, using only the fast electronic part of water dielectric constant. This approach is formerly equivalent to a rescaling of ionic charges by a factor 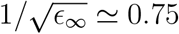^46^.^74^ Previous studies however suggested that a 0.8 scaling factor yielded better ion pairing behavior, in part due to the implicit inclusion of some electronic effects within the classically used water force fields,^49,55^ but also because the real scaling factor should slightly depend on the distance between ions—varying between about 0.85 at contact to 0.75 at large distances— so that the choice of a single uniform scaling factor is only an approximation.^75^ For monovalent cations and anions (K^+^, Na^+^, Cl^−^), ECC variants of the parent force fields were derived for each combination of ion and water force fields following the previously validated strategy:^50,57,76^ in addition to scaling the charge by a 0.8 factor, the Lennard-Jones radius of the ions, *σ*, was slightly reduced to recover proper solvatation properties, and in particular the position of the first peak of the ion-water radial distribution function (see parameters in bold on Table SI-1, typically decreasing *σ* by about 5-10%). For the phosphates, the charge rescaling was applied to the 4 atoms 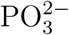, and the overall −1.6 charge was adjusted using the hydroxyl oxygen (parameters in orange on Figure 1), without any modification of the Lennard-Jones parameters.

**Figure 1:**
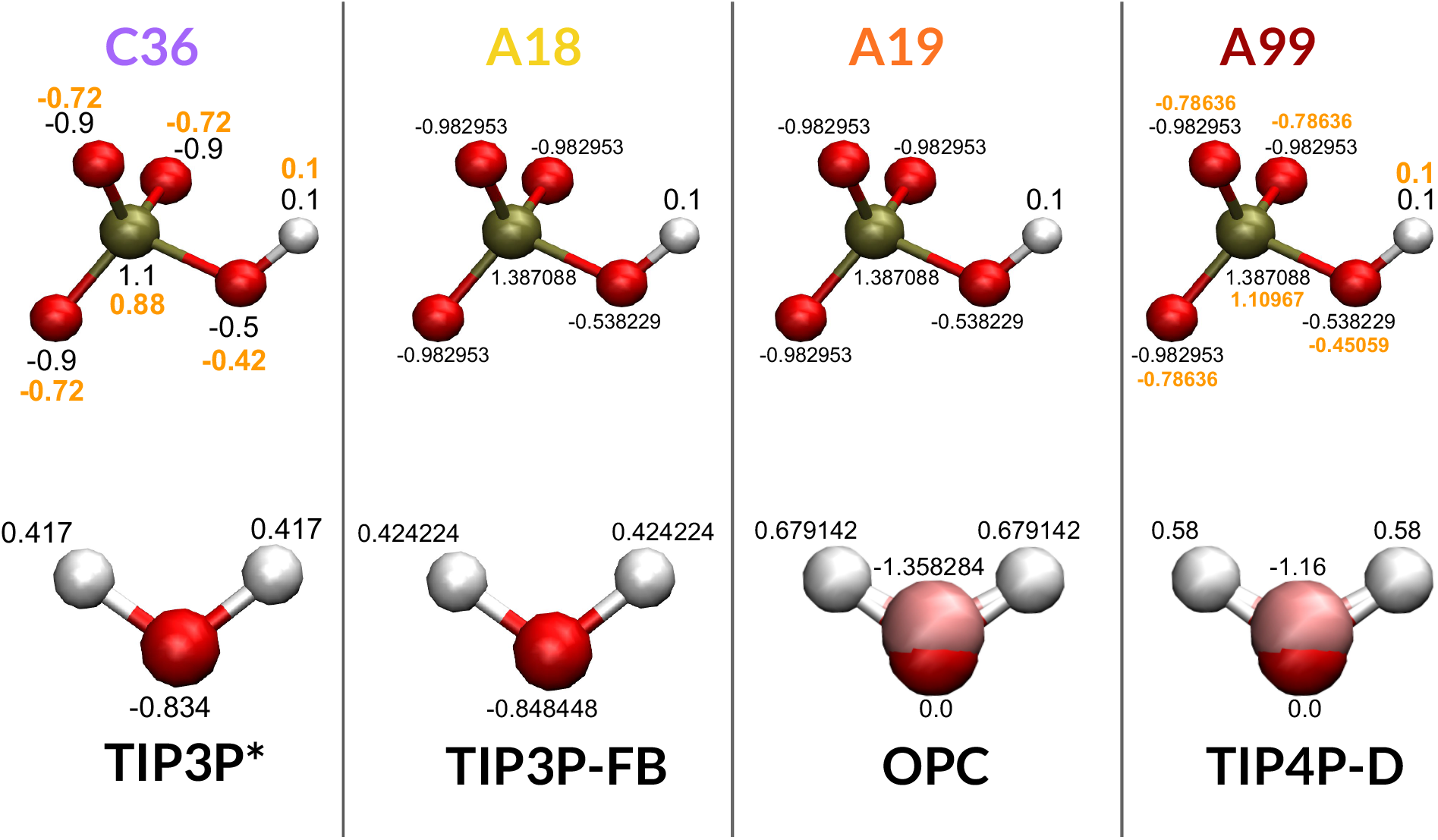
Partial charges for the phosphate molecule and water model of each forcefield. Full charges are in black, ECC-corrected charges are in orange. Water molecules are not to scale with the phosphates for visibility.

For the protein forcefield, rescaling was performed manually on the topology, using a very similar strategy to that successfully used in a recent paper on ion channels that came out during the preparation of this manuscript. ^53^ A first attempt, which we will call “ECC-IP”, is to rescale only the bulk ions and the phoshorylated groups. For phosphoresidues, the rescaling was performed similarly as for the phosphates (see Figures SI-1 and SI-2). Another attempt called here “ECC-IPP” is to rescale all the charged residues of the protein, including its termini. For the CHARMM force fields where the charge is localized, we applied the scaling factor exclusively on the charged group. For Amber force fields, we applied the charge scaling on a chemically intuitive group of atoms of the side chains. The charge on the previous side-chain carbon atom was then adjusted so as to obtain a 0.0, +0.8, −0.8 or −1.6 total charge depending on the amino acid type. Details of the employed ECC variants are provided in the SI (see Figures SI-1 to SI-6) and parameter files are provided in a public Zenodo repository : https://doi.org/10.5281/zenodo.16980659

We insist that these ECC variants of each force field are not finely optimized and should be regarded as a mere proof of concept regarding the impact of the ECC correction on ion pairing with phosphorylated IDP and IDP conformational ensemble.

### Calculation of osmotic coefficients

To assess the quality of ion pairing to the doubly negatively charged phosphorylated moiety in different force fields, we used 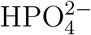 as a simple proxy for the phosphate group. While 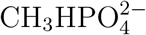 would have been a closer proxy for the phosphorylated side chain, no experimental osmotic coefficients are available in the literature, which led us to use 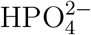 instead, where osmotic coefficient data is reported both for its K^+^ and Na^+^ salts.^77,78^ In each case, we built 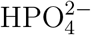 by analogy with the parent force field, keeping the same partial charges, Lennard-Jones, angle and dihedral parameters for the 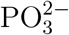 part of the compound as in the parent phosphoserine. The charge of the hydrogen was set to 0.1. The overall −2 charge of the phosphate was completed by tuning the partial charge of the remaining hydroxyl oxygen (see Figure 1).

For each force field, we first performed a steepest descent minimization for 50000 steps of the 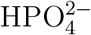 group in vaccuo using GROMACS v2021.5 in order to set the proper internal geometry.^79^ The cut-off scheme was defined as Verlet while the long range electrostatic interaction cutoff was extended to 1.5 nm in order to ensure that the entire compound would be comprised within it.

To compute the osmotic coefficient of 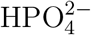 solutions across different force field variants, we employed a strategy inspired by the original protocol of Luo and Roux,^44^ adapted to a spherical confinement setup as described by Piana et al..^56^ 32 copies of 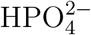 were randomly placed in a 12nm-side periodic box. After addition of water, 64 random water molecules were swapped for sodium or potassium ions to insure the neutrality of the box, resulting in a 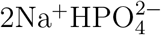 or a 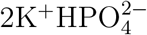 system. It was then minimized for 50000 steps treating the electrostatic interactions with a PME scheme and with short-range cutoffs set to 1 nm.

The four forcefields studied here each have their own requirements in terms of barostat, thermostat, and cutoffs. More details are given in Ref^29^ and in the provided parameters files. For all systems, the integration timestep was 2 fs, the temperature was set to 300 K and pressure to 1 bar, and simulations were ran with GROMACS v2022.4. We started the equilibration process by a 1 ns run in the NPT ensemble followed by 500 ps in the NVT ensemble with no restraints to relax the system. We used PLUMED^80^ v2022.4 to restrain the positions of the phosphates and cations at the center of the box within a sphere of radius 2.3 nm. The force constant *k* of the moving restraint was then gradually increased from 0 to 10 kcal.mol^*−*1^.A^*−*2^ over the course of 400 ps. The last frame of this step was further equilibrated with a 10 ns run in the NPT ensemble, with a spherical restraint radius *R* adapted to the desired concentration.

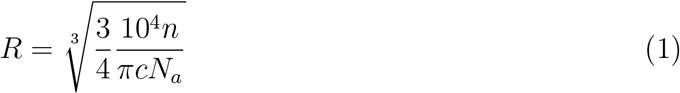

with *n* the number of phosphates, *c* the concentration in mol.L^*−*1^ and *N*_*a*_ the Avogadro number.

The osmotic coefficient *ϕ* is defined as *ϕ* = *P/P*_0_, where *P* is the osmotic pressure and *P*_0_ the perfect gas pressure. *P*_0_ can be expressed as *P*_0_ = *s*.*c*.*R*.*T* where *s* is the number of reactants (3 in our case, 2 cations and 1 phosphate), *c* is the molar concentration, *R* is the perfect gas constant and *T* is the temperature. The osmotic pressure can then be derived by calculating the average force ⟨*F* ⟩ = ⟨2*k*(*r* − *R*)⟩ exerted by the solute on the harmonic wall of area *A* = 4*πR*^2^, where *r* is the distance of a solute particle to the center of the box. *R* is equal to 3.988 nm, 2.938 nm and 2.512 nm for concentrations of 0.2, 0.5 and 0.8 mol.L^*−*1^ respectively. 10 replicas of 25 ns were run from the last frame of the resulting equilibration for each system. Frames were saved every 10 ps, and distances *r* from the center of the box to the phosphates and cations were saved with PLUMED every 1 ps to perform the average force calculation. Cluster numbers of 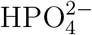 molecules were computed using the **gmx clustsize** function of GROMACS using a 0.6nm cutoff.

### Binding free energy calculations for binding of 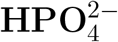 with monovalent cations

The standard binding free energy between a cation and the 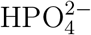 anion, 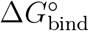, was computed for different forcefields using an alchemical simulation strategy^81–85^ within the double decoupling theoretical framework.^85–89^ Two alchemical transformations are performed: the cation is gradually decoupled from its environment either in bulk water 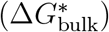 or within the ion pair 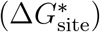, with flat-bottom harmonic restraints applied to maintain the cation in the desired configuration during the transformation. A correction term Δ*G*_PBC_ accounts for the net-charge artifacts introduced by periodic boundary conditions,^90^ and an additional correction 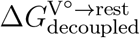 accounts for the application of restraints in the decoupled state, relative to the standard-state concentration.^55^ 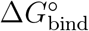 is then obtained by summing these separate contributions:

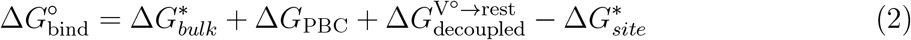

A detailed discussion of this framework is available in Refs.^91,92^ Contact and solvent-shared ion pair (SShIP) geometries were defined based on the first and second minima of the radial distribution function, g(r), between the cation and the P atom. The corresponding distance ranges are listed in Table SI-2. Alchemical simulations were performed using Gromacs v2023 in cubic boxes containing one 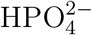 anion, one cation, and 905 or 896 water molecules for A99 and C36, respectively. Electrostatic interactions were first decoupled over 11 *λ*_elec_ windows (1.0 to 0.0, step 0.1), followed by van der Waals decoupling over 4 *λ*_vdw_ windows (0.7, 0.5, 0.1, 0.0). Each window was equilibrated for 250 ps and sampled for an additional 5 ns.

Free energies were calculated using the Bennett Acceptance Ratio (BAR) method^93^ as implemented in Gromacs. For each combination of forcefield and binding geometry, three independent alchemical replicates were performed. The reported 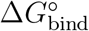 values correspond to the mean over these three replicates, and the error bars shown in the bar plots represent the standard deviation across them. Our protocol —with a careful choice of windows and restraints, as well as extensive sampling in each window— yields very robust free energy estimates, with standard deviations smaller than 1 kJ·mol^−1^ over 3 replicas.

### Molecular dynamics simulations of a highly phosphorylated peptide

We used the sequence and phosphorylation state of the rhodopsin 7PP peptide described in Ref.^94^ The peptide was built using the tleap program from AmberTools22 which generates a fully extended conformation.^95^ Phosphorylations were added using CHARMM-GUI.^96^ The phosphorylated peptide was then solvated in a box of sidelength 9 nm. The concentration was set to 0.1mol.L^*−*1^ to reproduce experimental conditions, resulting in the addition of 60 cations (Na^+^) and 43 anions (Cl^−^). For the ECC-IP peptide, as only some charged residues are scaled, the box is not overall neutral. We added another cation in order to bring the total charge of the box as close to 0 as possible (+0.2). The system was then minimized for 50000 steps. The minimized output was used to create 3 independent replicas. For A99, a first equilibration step of 200 ps was performed in the NPT ensemble to set the correct pressure in the waterbox, followed by a 2 ns run in the NVT ensemble with the peptide heavy atoms being restrained. For C36, only the 2 ns step was performed since the water box is already at the correct pressure. The production step lasted 520 ns per replica and frames were saved every 10 ps.

### Analyses

Analyses were performed using Python scripts based on the Numpy and MDAnalysis libraries.^97,98^ Plots were displayed with Matplotlib,^99^ and images were rendered using Visual Molecular Dynamics. ^100^ Local Curvatures (LCs) and Local Flexibilities (LFs) were computed using the MDAKit Menger_Curvature.^101,102^ Chemical shift (CS) predictions were performed with SPARTA+ (*H*_*N*_ and *H*_*α*_) and PPM (*H*_*N*_) every 10 frames before averaging over all collected values from the three replicas.^103,104^ CSs can be normalized in order to remove the resonance dependency per amino acid by subtracting the random coil value, giving the Secondary Chemical Shifts (SCSs).^105^ We extracted the CSs values for *H*_*N*_ and *H*_*α*_ from Kisselev et al^94^ and used the dataset from Kjaergaard et al for random coil values of canonical amino acids.^106^ Random coil values for phosphoresidues can vary based on the used reference peptide.^107^ For the phosphoserines and phosphothreonines, we thus chose the values reported in the most recent study by Hendus-Altenburger et al.^108^ A *n*P-collab was considered to be formed in a simulation if a cation was present within 4 Å of *n* phosphorus atoms.

## RESULTS

### Assessing the interaction between phosphorylated moieties and cations against osmotic coefficient data

To assess which force field combination provides the more realistic description of the interactions between phosphorylated moieties and cations, we used as a simple proxy 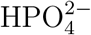, where osmotic coefficient measurements are reported in the literature over a range of concentrations for both the potassium and sodium salts.^77,78^ Osmotic coefficients of electrolytes quantify the tendency of ions to associate in solution and deviate from the ideal solution behavior,^109,110^ and are thus sensitive probes of ion pairing. They are often used to assess the quality of force fields to describe the structure of electrolytes, or even as targets for ion force field development.^52,56,110,111^ The closest chemical proxy to a phosphoserine would have been methylphosphate 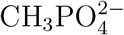. However, as far as we have been able to determine, no osmotic coefficient value has been reported for this compound, presumably due to chemical instability. Our strategy was thus to build for each force field an associated model for 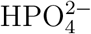, that keeps the same charge and atom types on the 4 key interacting atoms (PO_3_) as in the parent force field. These models are thus directly reflecting the ion pairing behavior of the corresponding phosphorylated moieties in the parent protein force field (see Methods).

For all the different force field combinations (C36, A19, A99, with different ion and water models, see Methods), we first computed the osmotic coefficient *ϕ* of sodium and potassium 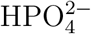 electrolytes, at a concentration of 0.2 mol.L^*−*1^, close to the ionic physiological concentration (Figure 2 and Table SI-3). Experimental osmotic coefficients are very similar for Na^+^ and K^+^, for both experimental sets of measurements (respectively 0.7504 and 0.7620 in the dataset by El Guendouzi and Benbiyi ^77^ and 0.753 and 0.767 in that by Goldberg ^78^). All tested forcefield combinations underestimate the osmotic coefficients, more dramatically for Na^+^ than K^+^. Since a lower osmotic coefficient is associated with stronger interactions between the solute components, this means that the cation/phosphate interaction in all forcefields is (sometimes severely) overestimated. For A19, the osmotic coefficients for both Na^+^ and K^+^ are relatively close to the experimental values. A99 performs a little worse, with the K^+^ coefficient being three quarters and that for Na^+^ being half of the experimental values. A18 has the worst results out of the Amber family with a halved value for the K^+^ salt with respect to experiments and an extremely low coefficient for the Na^+^ solution (0.11 ± 0.02). C36 exhibits the largest discrepancy between Na^+^ and K^+^ (0.11 ± 0.02 vs 0.68 ± 0.02). While the K^+^ coefficient of C36 is the best out of the four forcefields, its Na^+^ coefficient is comparatively as bad as with A18.

**Figure 2:**
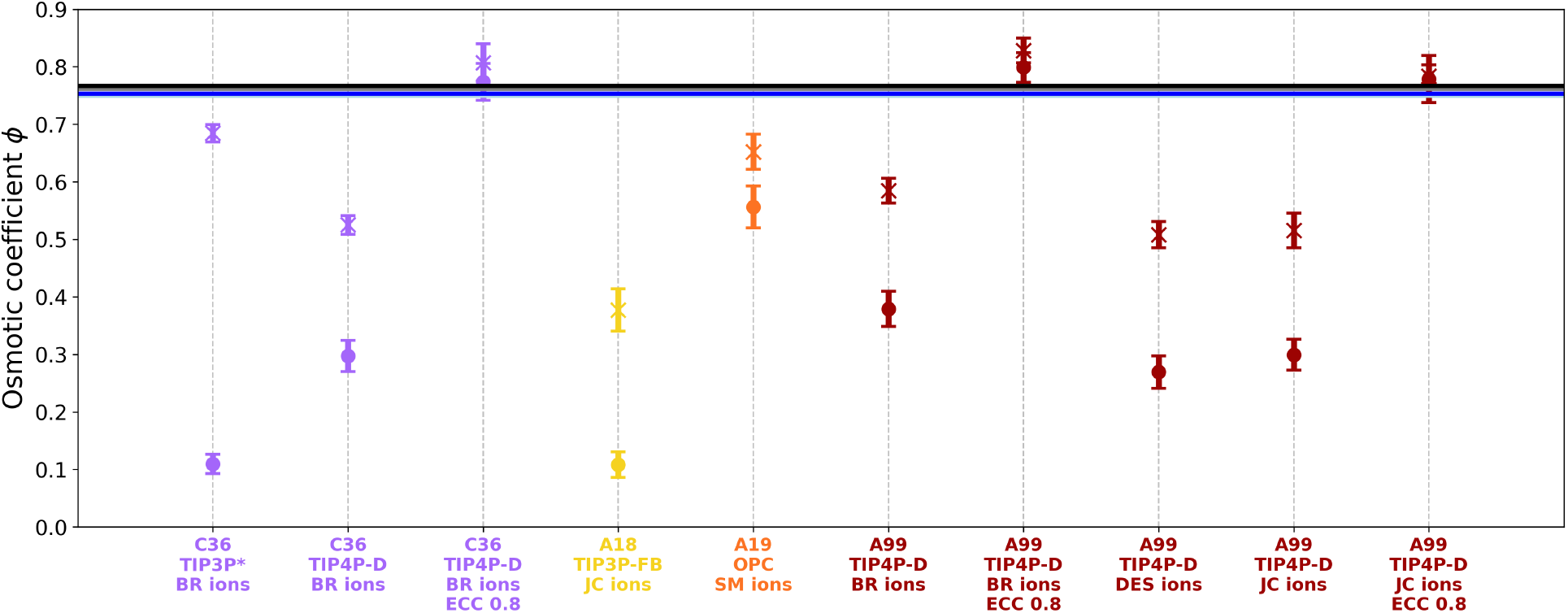
Computed osmotic coefficients at concentration 0.2mol.L^*−*1^ for C36 (purple), A18 (yellow), A19 (orange) and A99 (red). Coefficients for Na^+^ and K^+^ are represented with a round and cross shape, respectively. Experimental values measured by El Guendouzi and Benbiyi ^77^ are represented by a dark blue line at 0.7504 and a black line at 0.7620 for Na^+^ and K^+^ respectively. Experimental values measured by Goldberg ^78^ are represented by a light blue line at 0.753 and a grey line at 0.767 for Na^+^ and K^+^ respectively. Standard deviation is computed over 10 replicas.

These results can be interpreted under the light of the earlier study by some of us on the formation of cation-mediated bridges between phosphorylated residues, what we called the *n*P-collab phenomenon. ^29^ We simulated a tri-phosphorylated peptide which showed different levels of interactions mediated by counterion bridges depending on the forcefield. The level of *n*P-collabs with A19 was very low, with very transient interactions between phosphoserines and cations, coherent with the higher values of *ϕ* reported on Figure 2. The rest of the results are also coherent overall with the intensity of *n*P-collabs that were previously determined, for example with C36 and A18 phosphoresidues forming highly stable bridges through Na^+^ and A99 forming only transient bridges, in accordance with the low and high *ϕ* coefficients of the forcefields respectively.

Overall, these results reveal a marked overbinding of cations by phosphorylated moieties in all common non-polarizable force fields, that probably leads to an overestimation of the number and lifetime of cation-mediated bridges between phosphorylated moieties. There is thus an accute need for improving the ion pairing behavior of phosphorylated aminoacids to obtain a more realistic behavior of phosphorylated IDPs in molecular simulations.

### Improving the ion pairing behavior of phosphorylations

We decided to focus our efforts to improve the interaction strength on C36 and A99 since we showed previously that A18 and A19 both have issues in their parameterization of phosphorylations which are independent from the interactions with cations.^29^ We tested several strategies to try to improve the osmotic coefficient values.

Since A19 and A99 perform best with 4-point water models, we hypothesize that using one of them for C36 might be key for improvement. We then repeated our simulations with the TIP4P-D water model.^112^ While the osmotic coefficient of the Na^+^ is improved but still very far from the experimental reference, that for K^+^ is degraded. This suggests that while the water model plays a key role in the cation/phosphate interaction strength, the latter is the resultant of a subtle equilibrium between many parameters so that a change of water model does not always lead to consistent improvements for both salts.

We next decided to investigate how sensitive to the choice of ionic parameters was the osmotic coefficient. We thus repeated the osmotic coefficient calculation for A99, replacing the cation parameters by those prescribed in the DES-Amber SF1.0 forcefield ^56^ or those from Joung and Cheatham (abbreviated as JC).^61^ Parameters from the latter perform worse than the original A99, and parameters from DES-Amber SF1.0 even more so. This seems to imply that an individual parameter set is not to blame, but that the problem is instead common to all non-polarizable full-charge force fields, quite independently of both the water model and the ion parameter set.

We then turned to testing the ECC strategy, which implicitly takes into account electronic polarization by scaling the charges of ions and charged biomolecular groups, as a way to improve systematically all ionic interactions in our simulations. Following previous work, we used a 0.8 scaling factor for both monovalent ions and, for a start, the 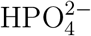 molecularion (see Methods). Considering that changing the water model for C36 from TIP3P* to TIP4P-D presented a noticeable improvement for Na^+^, we decided to keep working with TIP4P-D for the ECC correction (called ECC-C36 in this part). The corrected results for Na^+^ (0.78 ± 0.03) and K^+^ (0.81 ± 0.03) are in remarkable agreement with the experimental values. We next calculated *ϕ* for A99 corrected with the ECC, deriving ECC0.8 variants of both the BR and JC ion parameter sets. Both variants strongly improve the osmotic coefficient compared to the experimental values for both cations, even if the BR ECC0.8 variant slightly overestimates *ϕ* for both Na^+^ (0.80 ± 0.02) and K^+^ (0.83 ± 0.02). The JC ECC0.8 variant performs stunningly for both Na^+^ (0.78±0.04) and K^+^ (0.78±0.02). We thus decided to select this variant as our best ECC-corrected forcefield for A99 (called ECC-A99 in this part).

To assess whether the ECC correction would still hold in more concentrated solutions, additional calculations of *ϕ* were performed at concentrations 0.5 and 0.8 mol.L^*−*1^ (Figure 3 and Tables SI-4 and SI-5). We first studied how the regular full charge forcefields C36 and A99 behaved at higher concentrations. In line with the discrepancy at 0.2 mol.L^*−*1^, Na^+^ in C36 displays *ϕ* values much lower than experiments. More surprisingly, *ϕ* values for K^+^ degrade a lot compared to experiment as concentration is increased. The opposite trend is observed for A99, with *ϕ* values of Na^+^ decreasing too quickly with increasing concentration, while values for K^+^ decrease at a similar rate as the experiment. Overall, the original full charge forcefields fail to reproduce the experimental measurements both in trend and accuracy.

**Figure 3:**
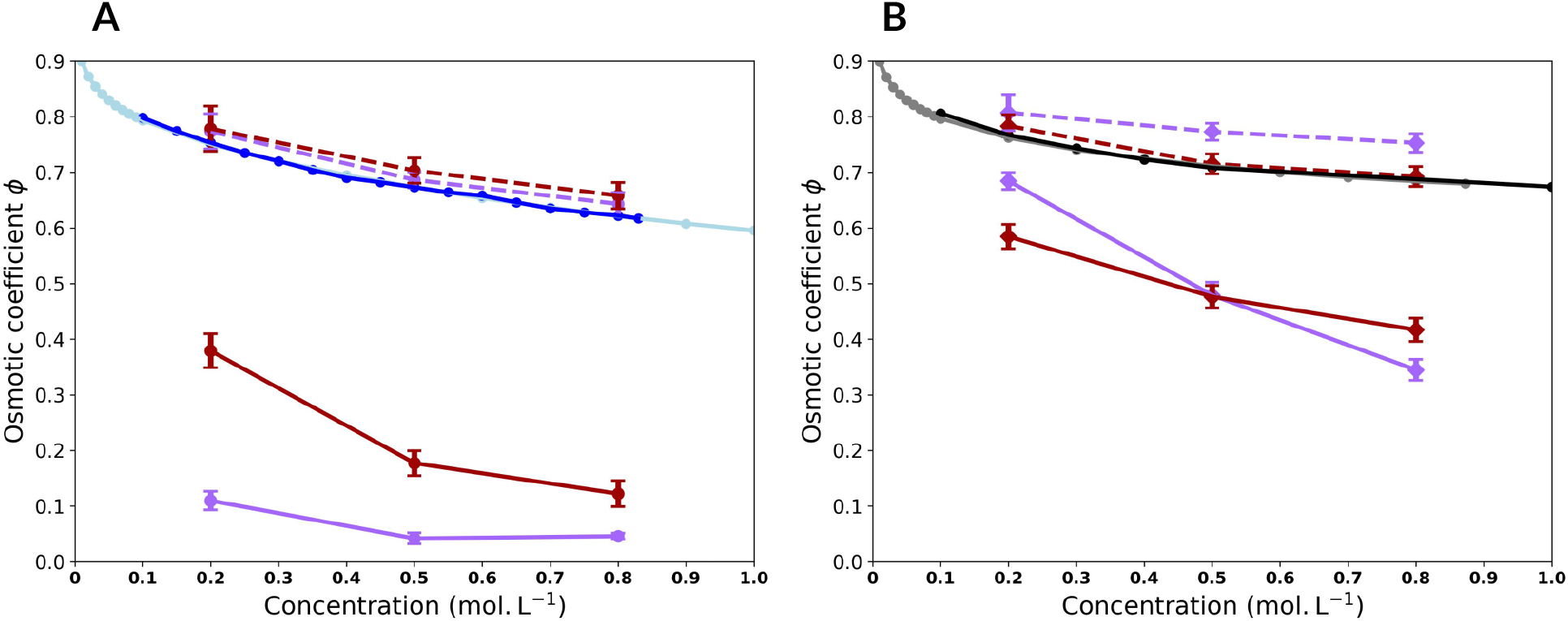
Osmotic coefficients for C36 (purple) and A99 (red) for the classical forcefield (full line) and the ECC-rescaled forcefield (dotted line) as a function of the concentration. A) Na^+^, B) K^+^. Experimental values measured by El Guedouzi et al^77^ are represented by a dark blue line and a black line for Na^+^ and K^+^ respectively. Experimental values measured by Goldberg et al^78^ are represented by a light blue line and a grey line for Na^+^ and K^+^ respectively. Standard deviation is computed over 10 replicas.

We then assessed how the ECC force field variants perform at these higher concentrations. For both corrected forcefields, the agreement in trend and accuracy for the Na^+^ solutions is remarkable. ECC-C36 follows closely the experimental curve and so does ECC-A99 despite a slight overestimation compared to ECC-C36. For K^+^, this behavior is inverted, with ECC-A99 perfectly following the experimental curve and ECC-C36 mildly overestimating *ϕ* at higher concentrations.

Overall these results point to a remarkable improvement of ion pairing with our phosphate proxy 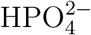 by using the ECC charge scaling strategy. The ECC variants manage to restore the agreement between the computationnaly-derived and the experimental osmotic coefficients for a variety of forcefields, cation types and concentrations. The ECC correction also elicits significant changes in the aggregation behavior of the solute. At 0.2 mol.L^*−*1^, while C36 phosphates and Na^+^ cations tend to form large clusters by the end of the simulation, with ECC-C36 the solute remains mostly homogeneous in the solvent (Figure SI-7). In addition, we explicitly computed the number of clusters formed by the 32 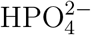 molecules (which can be interpreted as aggregates mediated by the bulk cations) at 0.2 mol.L^*−*1^ for the regular forcefields and the ECC-corrected ones (Figure SI-8). As could be expected, based on their osmotic coefficient, 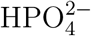 molecules with A18 and C36 heavily aggregate in the presence of Na^+^, A99 slightly less and A19 almost does not, while molecules remain largely unaggregated in the presence of K^+^ for all forcefields. The application of the ECC correction to A99 and C36 completely modifies the interaction profile of the 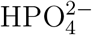 molecules, as more than 80% of the conformations display no interaction at all, and the 20 remaining percent consist of one interaction or two at most. This modification also impacts the distribution of water around the phosphate molecules (Figure SI-9), leading to the restoration of a defined first water shell for C36 in sodium, and to a larger radius of the shell for both ECC-C36 and ECC-A99 in both cations.

This tendency of the regular forcefield for phosphates and cations to cluster is in line with the observed formation of multiple cation bridges (*n*P-collabs) in simulation of phosphorylated peptides with full charge force fields,^29^ and its removal by the ECC correction could mean that the number and lifetime of *n*P-collabs observed with common non-polarizable force fields might be strongly overestimated.

In light of the strong improvement in ion pairing brought about by the ECC variants, we sought an in-depth understanding at the molecular level of the changes in ion pairing linked to the ECC variants. To this aim, we examined the relative stability of the contact and solvent-shared (separated by a single water molecule) ion pairs with both the original A99 and C36 force fields, and their ECC variants. We thus computed for these various force fields the binding free energy (Δ*G*) between 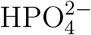 and either Na^+^ or K^+^ (see Figure 4 and Tables SI-6 and SI-7 for the values in each replica). Note that no experimental 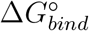 values are available for these specific ion pairs, so these calculations are only used to investigate in details the changes in ion pairing behavior underlying the previously observed changes in osmotic coefficients. Binding free energies with 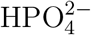 are only reported for divalent ions^113^ (−16.2 kJ·mol^−1^ and −15.5 kJ·mol^−1^ for Mg^2+^ and Ca^2+^, respectively) which are, as expected, significantly stronger than the computed binding with monovalent ions for all forcefields.

**Figure 4:**
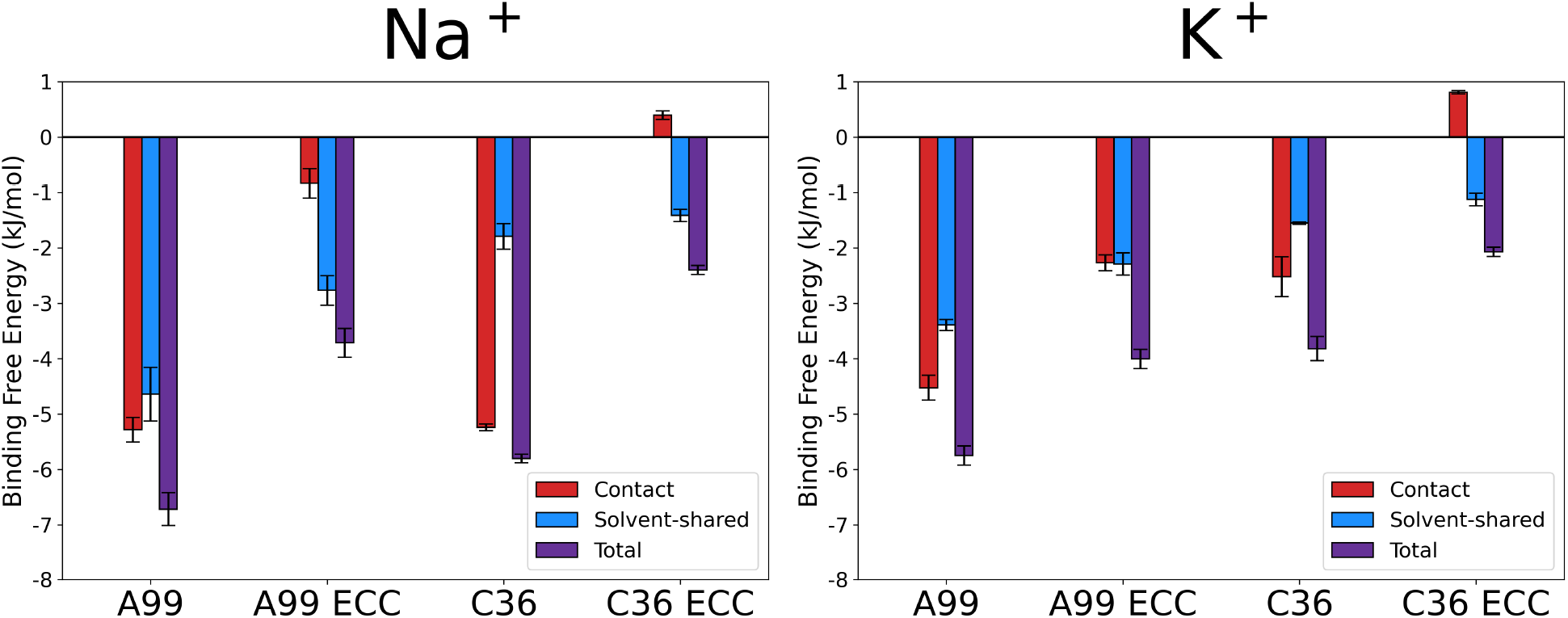
Standard binding free energy (Δ*G*^°^) between 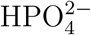 and a) Na^+^ or b) K^+^. The total energy (in purple) was decomposed in its direct contact component (in red) and the solvent-shared (SShIP) component (in blue). Standard deviations were calculated on three replicas.

As expected, ECC variants systematically reduce the overall binding free energy, consistently with the previously described increase in osmotic coefficient, which also points to a decreased formation of ion pairs (Figure 2). For both force fields (A99 or C36), the decrease in binding free energy after the ECC correction 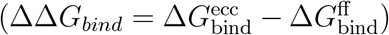 is more pronounced for Na^+^ than K^+^ (for instance for A99, ΔΔ*G* = 3.0 kJ·mol^−1^ for Na^+^ but only ΔΔ*G* = 1.7 kJ·mol^−1^ for K^+^).

These free energy calculations also allow us to examine how different force field variants change the relative stability of different types of ion pairs. Focusing on A99-ECC, which yielded for K^+^ the osmotic coefficients closest to experiment, the ECC correction simultaneously reduces the stability of both contact (by 2.2 kJ·mol^−1^) and solvent-shared ion pairs (by 1.0 kJ·mol^−1^) compared to the parent full-charge A99 force field. With the A99-ECC forcefield, both ion pair geometries are found to be very similar in terms of free energy. However, we limit our study here to a 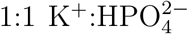 stoichiometry, while realistic conditions may involve higher cation:phosphate ratios (e.g. 2:1), which could modulate the stability and distribution of binding modes. In contrast, C36-ECC yields much weaker ion pairs, with the contact pairing with both cations associated with a positive binding free energy (0.3 kJ·mol^−1^ and 0.8 kJ·mol^−1^ for Na^+^ and K^+^ respectively), and a very weak total Δ*G*^°^ (−2.4 kJ·mol^−1^ and −2.1 kJ·mol^−1^ for Na^+^ and K^+^ respectively) dominated by the solvent-shared ion pair. This result is in line with the overestimated osmotic coefficient observed for this forcefield, which suggests that ion pairing is underestimated by C36-ECC.

A general trend across these different forcefields is that while the full-charge forcefields predict the contact ion pair to be the most stable —sometimes strongly dominating the overall binding free energy, as for C36 and Na^+^— all ECC variants shift this equilibrium in favor of the solvent-shared ion pair, even if both ion pair geometries are very similar in free energy for K^+^ of A99-ECC. This should then decrease the number of long-lived cation-mediated bridges between phosphorylated residues (*n*P-collabs) which rely on contact ion pairs.

While our computed binding free energies are often qualitatively consistent with trends in osmotic coefficients (stronger binding free energies and increased ion pairing being associated to weaker osmotic coefficient), our results show that two forcefields with similar binding free energies can be associated with very different osmotic coefficients (Na^+^ of A99-ECC and C36). This indicates that 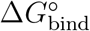 and osmotic coefficients probe distinct aspects of ion–phosphate interactions. While Δ*G*^°^ reflects a local, pairwise tendency to form ion pairs, *ϕ* is sensitive to collective organization and long-range ion clustering, which can differ significantly between forcefields even when local binding appears comparable. Those two physical quantities are thus complementary to assess ion pairing behavior. Unfortunately, there is currently no experimental technique capable of directly quantifying the relative populations of contact versus solvent-shared phosphate binding modes in these systems. This lack of data constitutes a major limitation for validating simulations at this level of resolution—it is thus impossible to assess whether the very low stability of the contact ion pair with C36-ECC is physically relevant or not.

### Testing the ECC parameters on an experimentally-characterized peptide with multiple phosphorylations

In order to test whether the ECC-corrected phosphates and cations better represent the conformational ensemble of a multiphosphorylated peptide we attempted to find NMR data of polyphosphorylated peptides. Unfortunately, the acquisition of a NMR spectra is usually performed at very low salt concentration, and studying multiple phosphorylations on a peptide is rarer than single-point phosphorylation. We only identified one work with a significant concentration of 0.1 M sodium phosphate buffer, the study of the C-terminal tail of rhodopsin (residues 330-348) by Kisselev et al. in 2004.^94^ The study notably provides proton resonance data that we will later attempt to reproduce. Although a strong assumption, we considered that the buffer could be approximated by a simple NaCl concentration. The 7PP peptide is composed of 19 residues of sequence DDEASTTVSKTETSQVAPA. The three serines and four threonines are phosphorylated, increasing the total charge of the molecule from −4 to −18 since phosphorylations are in their fully deteriorated form at pH=7,^33^ hence the name 7PP of the peptide. The peptide was confirmed to be fully disordered by the measurements and expected to be rather extended, in agreement with an earlier study by Langen et al. using a spin-labeling strategy.^114^

We first attempted to reproduce the global properties of the peptide in sodium chloride using molecular dynamics simulations with the original forcefields C36 and A99, then with their corrected version with the ECC correction applied solely on the phosphate groups of the phosphoresidues and on the Na^+^ cations (ECC-IP). We then tested a variant where all the charged groups of the peptides are modified by charge scaling (ECC-IPP, see Methods). We started with computing the radius of gyration *R*_*g*_ and the end-to-end distance *R*_*ee*_ distributions for the concatenated three replicas with each forcefield (Figure 5). The regular C36 forcefield displays a very narrow distribution of *R*_*g*_ centered on 10 Å, and a distribution of *R*_*ee*_ centered on 25 Å, while both the ECC-IP and ECC-IPP forcefields see the average of their *R*_*g*_ distribution be shifted towards much higher values and a higher spread. The *R*_*ee*_ distribution is also shifted to much higher values, with the average end-to-end distance doubled. On the other hand, the *R*_*g*_ distribution of A99 is wider, spanning 6 Å and centered on 12 Å with what could be interpreted as two maxima. This is not the case for the ECC-corrected forcefields which only have one maxima around 12 Å. The same behavior is observed for *R*_*ee*_, with 2 maxima for A99 and only one maximum for the ECC-corrected forcefields (see Figure 5). The drastic change in the profiles of the distributions for C36 upon ECC correction is a clear sign that the peptide moved from a compact conformation to an extended one. The effect is less pronounced for A99 as it seems that the peptide was already quite extended before correction, but still present. This difference between the forcefields is in agreement with the behavior of the R2 peptide simulated in our previous work.^29^

**Figure 5:**
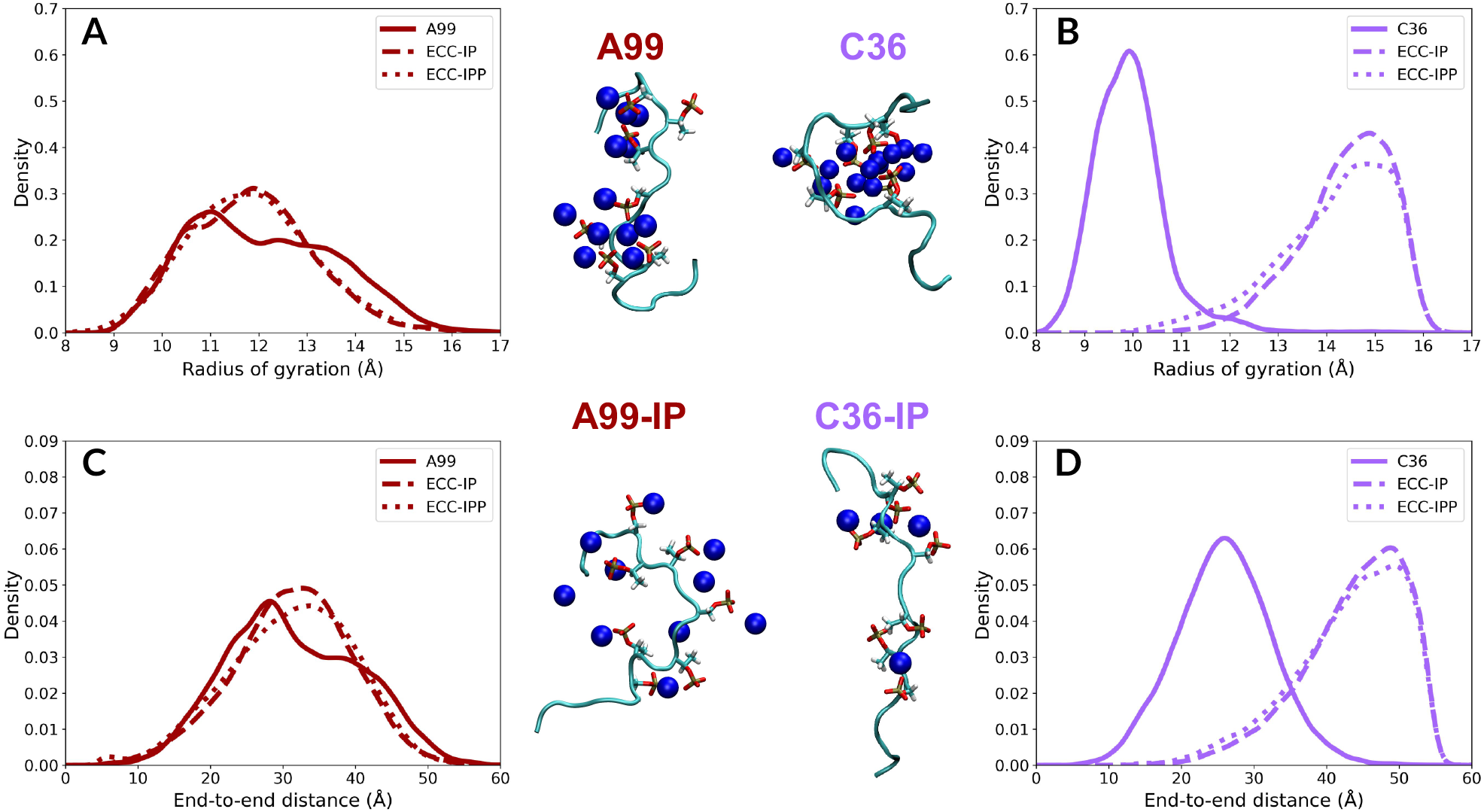
Distributions of the radius of gyration Rg of peptide 7PP for A99 (A) and C36 (B), and of the end-to-end distance Ree for A99 (C) and C36 (D). For the snapshots, the protein backbone is represented in cyan NewCartoon, the phosphoresidues in licorice, and the Na^+^ cations as dark blue Van der Waals spheres.

We next computed the Ramachandran plots of every amino-acid of the peptide for each forcefield (Figures SI-10 to SI-15) in order to assess the differences in backbone dynamics. A99 mostly explores regions related to beta-sheets and alpha-helices, while C36 samples in the PPII region. The ECC correction moves A99 towards more alpha-helicity, while for C36 it fully moves the backbone in the PPII region. Φ*/*Ψ distributions of the phosphoserine and phosphothreonine residues are without surprise most affected by the ECC, but residues comprised between them are also impacted. K339 for instance sees its backbone mostly sample the *α*-helix region for the full-charge C36 forcefield, but only samples the PPII region once the ECC is applied. Using the ECC correction thus not only improves the interaction with cations, but also has a significant impact on the backbone dynamics of the peptide (probably in part through the interaction with ions). Validating this behavior against experimental data is thus crucial, as we discuss in the manuscript, and attempted to do using pre-existing experimental Rg and NMR data.

Following our previous observations, we hypothesized that the difference of compaction between the regular forcefields and their ECC-corrected version lies in the number and strength of the cation-mediated bridges between phosphoresidues (*n*P-collabs). This is confirmed by the average number of potassium cations per frame involved in *n*P-collabs (bridges between *n* phosphoresidues) for each simulation (Figure 6), as both uncorrected forcefields present much more 2P-collabs than the ECC-corrected ones. For C36, 2P-collabs are 10 times more present than for the ECC-corrected forcefields, and it even exhibits 3P-collabs and 4P-collabs in significant amounts. For both the ECC-IP and ECC-IPP forcefields, 3P-collabs and more simply do not occur any longer. Interestingly, the number of direct ion pairs between a cation and a single phosphate group (mono-coordinated) is not impacted by the correction and remains around 2 cations per frame. On the contrary, for A99 the average number of mono-coordinated cations is nearly 5, almost as frequent as 2P-collabs for C36. This suggests that, although A99 has half the amount of 2P-collabs of C36, there is still a strong interaction between the phosphates and the sodium cations, but this interaction does not form bridges as easily. This is further implied by the small amount of 3P-collabs measured with A99 and the total absence of 4P-collabs. A99 ECC-IP and ECC-IPP reduce the phosphate/cation interaction as well, with a nearly halved amount of mono-coordinated cations, three times less 2P-collabs and a quasi-suppression of 3P-collabs. The amount of mono-coordinated cations actually reaches the same level as for C36, suggesting that a similar equilibrium is found for the interaction.

**Figure 6:**
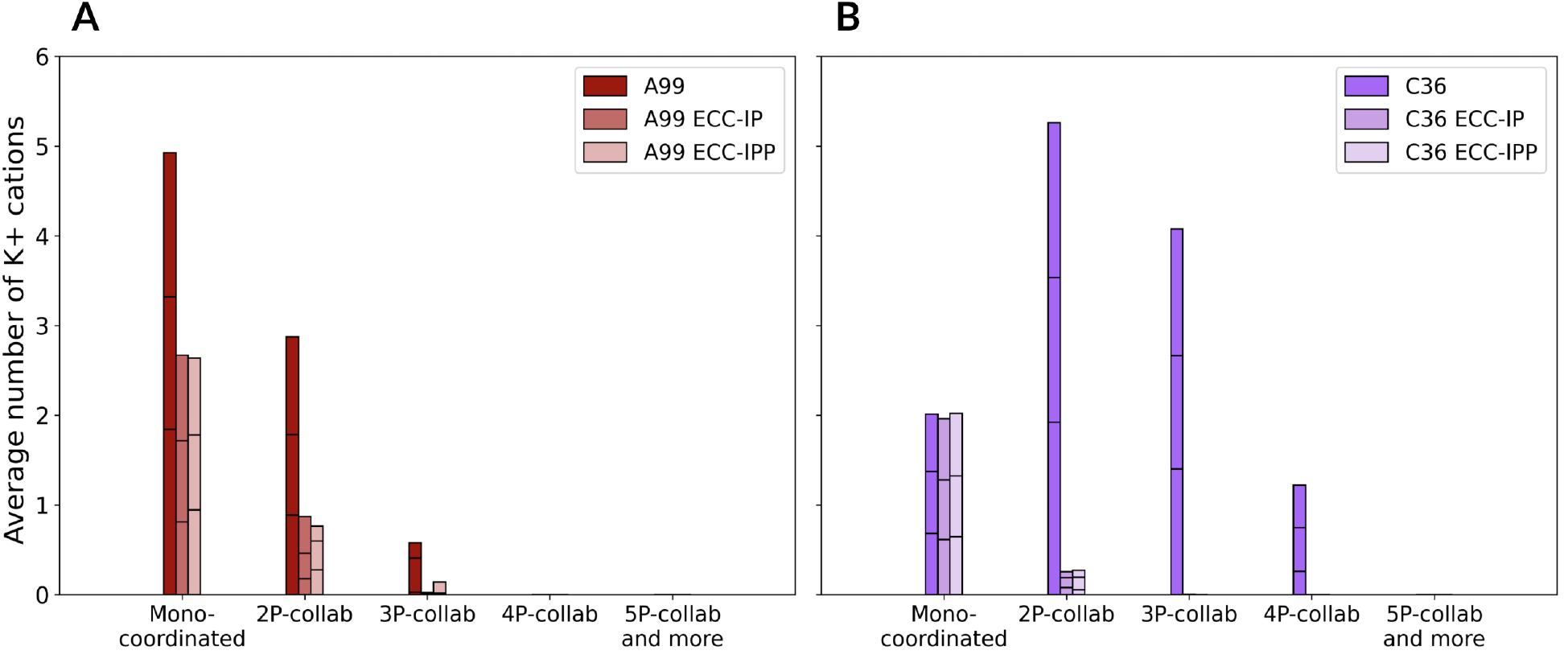
Average number of K^+^ cations per frame involved in mono-coordinated and *n*P-collabs with phosphorylated residues for A) C36 and ECC-rescaled forcefields, and for B) A99 and ECC-rescaled forcefields. The contribution of each replica is visualized by a black limit on the total bar. Simulations boxes contained a peptide with 7 phosphoresidues and 60, 61 and 60 cations for the full-charge, ECC-IP and ECC-IPP forcefields respectively.

To gain a better understanding of the dynamics of the interaction between the phosphoresidues and Na^+^ cations, the presence of the latter was checked near phosphates as a function of time (Figures SI-16 and SI-17). The profiles differ drastically between the original full charge and ECC-corrected forcefields. Sodium cations in both A99 and C36 tend to engage in long-lasting contacts with one or more phoshoresidues. These contacts can last up to 200 ns for A99 and lead to 2P- or 3P-collabs, and for C36 some cations can be stabilized in *n*P-collabs for the entire 520ns of the simulation, with several 3P- and 4P-collabs simultaneously present. The ECC correction completely modifies the timescale and pattern of the cations/phosphoresidues interactions. For the C36 corrected forcefields, contacts have a lifetime of around 20 ns, with a higher exchange between cations, and 2P-collabs are only maintained for 10 ns at most. For the A99 corrected forcefields, contacts can last longer (up to 50 ns) mostly in the form of 2P-collabs. Interestingly, the second replica of the A99 ECC-IPP is the only corrected replica presenting 3P-collabs.

We next conducted a detailed investigation of the specific phosphorylation pairs that interact via 2P-collabs (Figure 7). We observe that the original forcefields tend to have a more diverse and stable network of 2P-collabs than the ECC-corrected forcefields. Indeed, for the latter the 2P-collabs are mostly present for neighbor phosphoresidues (P6 with P7 or P13 with P14), while A99 and C36 can form 2P-collabs more often on longer sequence distances (for example P7 with P9, P9 with P11, P11 with P14). Interestingly, the ECC forcefield variants for A99 converge towards a similar 2P-collab pattern and intensity, with transient interactions happening between P6 and P7, and P13 and P14. On the other hand, C36 ECC-IP and ECC-IPP do not display the same pattern. This could imply that the additional correction on the charged residues and termini of C36 for ECC-IPP have an impact on the dynamics that is not felt by A99 ECC-IPP.

**Figure 7:**
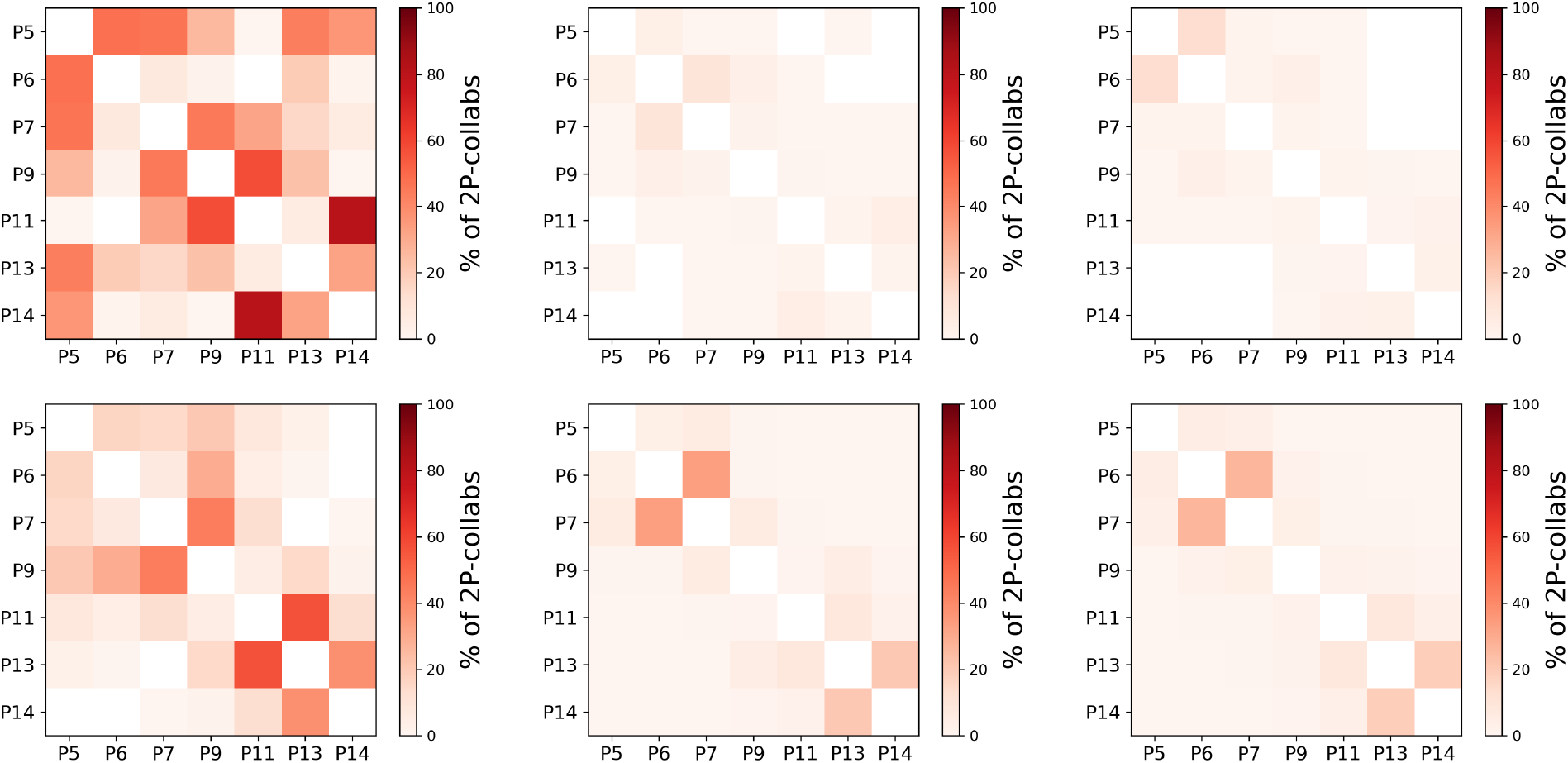
2P-collab percentage over the 3 replicas between the phosphoresidues of the 7PP peptide for the different forcefields. First row is C36, second row is A99. First column is full-charge forcefields, second column ECC-IP and third column ECC-IPP.

Finally, in order to probe how the dynamics of the backbone is modified, we employed the Local Curvatures (LCs) and Local Flexibilities (LFs) metrics (see Analysis section of the Material and methods).^102^ LCs represent the average curvature at a given residue and LFs the range of curvatures, which can be interpreted as a measure of flexibility. As could be expected, the curvature profile of the peptide simulated with the original C36 is very disparate both within the sequence and between the replicas (see Figure SI-18). For the ECC-corrected simulations however, the profiles converge and are flatter, which coupled to a high LF is usually indicative of a highly dynamic disordered behavior similar to a polyalanine.^29^ The same is observed for A99 and affiliated. From the LFs profiles, one can clearly measure the rigidity of the backbone induced by cation-mediated bridges (see Figure SI-19). Indeed, for C36, residues between the phosphorylation sites experience a drop from 0.05 Å^−1^ to below 0.02 Å^−1^. As a comparison, residues in a polyalanine have a LF of 0.07Å^−1^, while in a rigid globular core they are usually below 0.02 Å^−1^.^102^ The peptide is therefore fully constrained in its overall dynamics. A99 does not behave in the same way, with a wide variety of LFs profiles between the replicas, but some residues near the phosphorylations still experience very small LFs values. For both C36 and A99, introducing the ECC corrections does allow for an increased flexibility along the sequence. We also noticed that a minimum in LFs and LCs is usually reached for K10, which could indicate that this residue acts as a rigid hinge between the two halves of the peptide, and which could be a reason why this lysine is placed there among all these negatively charged residues.

All these results suggest that while the sodium/phosphate interaction for C36 is stronger and leads to more compact conformations, the ECC correction intensely reduces it, leading to extended conformations in much improved agreement with experiment. On the other hand, the sodium/phosphate interaction for A99 is not as strong to begin with, but the ECC correction does not weaken it as much, conserving the possibility for 2P-collabs to form and therefore allowing more compact conformations to still exist.

### Quantitative characterization at the residue level

A good descriptor of local dynamics for disordered proteins and peptides is the secondary chemical shifts (SCSs) since they are very sensitive to their chemical environment and its variations.^105^ The SPARTA+ program allowed us to compute both *H*_*N*_ and *H*_*α*_ SCSs (Figure 8), which we compare to the experimental reference^94^ (Figure 8, Tables 1, 2 and Tables SI-9 and SI-10). We will mainly focus on the relative performance between the forcefields rather than their ability to accurately reproduce the experimental signal. In particular, the capacity of each forcefield to reproduce the trend will be estimated by calculating the Pearson’s correlation coefficient *r* with the experimental data.

**Table 1:** RMSE and Pearson’s correlation coefficients *r* between the forcefield predictions of C36, C36+TIP4P-D, C36-IP and C36-IPP, and the experimental values for the SCSs of peptide 7PP with the SPARTA+ predictor. Best values are highlighted in bold.

| Forcefields | RMSE[H<sub>N</sub>] (ppm) | $r$[H<sub>N</sub>] (ppm) | RMSE[H<sub>α</sub>] (ppm) | $r$[H<sub>α</sub>] (ppm) |
| --- | --- | --- | --- | --- |
| C36 | 0.48 | 0.75 | 0.11 | 0.34 |
| C36+TIP4P-D | **0.34** | 0.87 | 0.10 | 0.44 |
| C36-IP | **0.34** | **0.89** | **0.08** | **0.65** |
| C36-IPP | 0.36 | 0.88 | **0.08** | 0.63 |

**Table 2:**
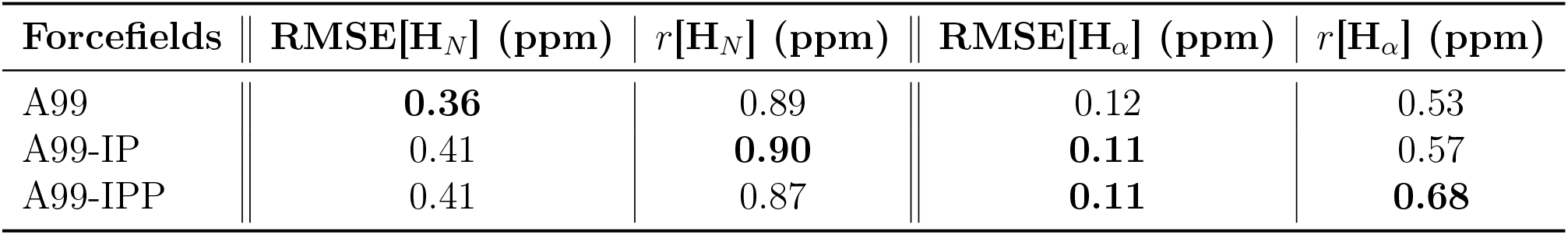
RMSE and Pearson’s correlation coefficients *r* between the forcefield predictions of A99, A99-IP and A99-IPP, and the experimental values for the SCSs of peptide 7PP with the SPARTA+ predictor. Best values are highlighted in bold.

**Figure 8:**
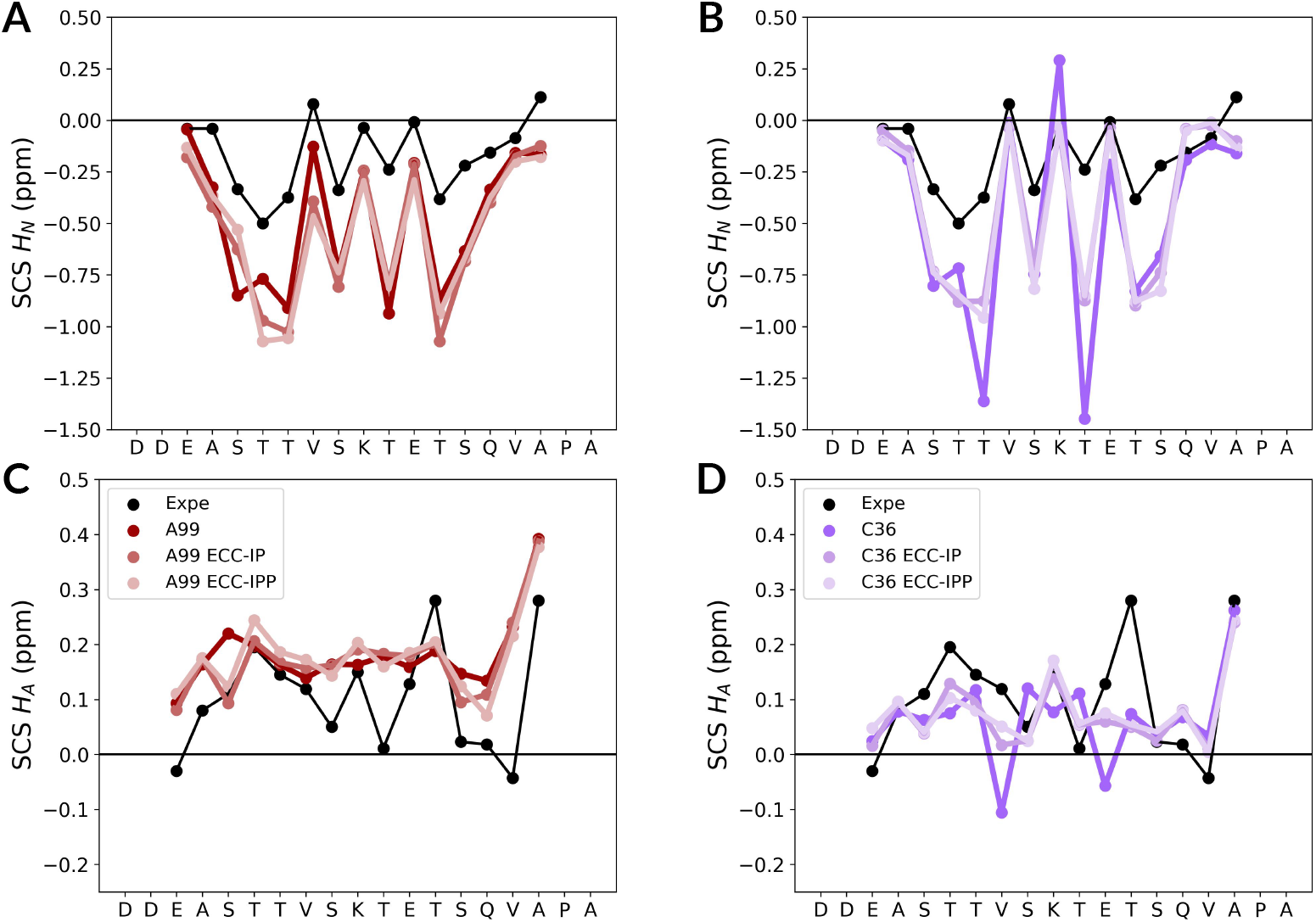
Secondary Chemical Shifts of *H*_*N*_ (upper row) and *H*_*A*_ (lower row) from the experimental data (in black) against computation on the 7PP peptide simulations using SPARTA+, excluding the first and last two residues of the 7PP sequence. *H*_*N*_ predictions for A) A99 and B) C36. *H*_*A*_ predictions for C) A99 and D) C36.

For the SCSs of *H*_*N*_, all A99-affiliated simulations underestimate the values. Almost all C36-derived results do the same, for both *H*_*N*_ and *H*_*α*_, while A99 simulations tend to overestimate the SCSs for the latter. The correlation of the signal for A99 is improved with the ECC-corrected forcefields, although no clear-cut winner can be crowned between A99-IP and A99-IPP. For the CHARMM family however, C36-IPP stands out as it improves all metrics compared to the other members and offers comparative performances to the A99-corrected forcefields. Considering that changing the water model from TIP3P* to TIP4P-D for C36 had a significant impact on the osmotic coefficient (Figure 2), we wondered how important the effect would be on the SCSs. We thus simulated 7PP with C36 and TIP4P-D without any ECC correction and computed the SCSs. In all cases, changing the water model improved the results, but not as greatly as the C36 ECC-IP or C36-IPP. The number of *n*P-collabs is reduced, especially 3P- and 4P-collabs, and the 2P-collab pattern by phosphoresidue pairs is slightly altered compared to the original C36 (Figure SI-20), but not to the extent of the ECC-corrected forcefields. Using a more advanced water model can increase the accuracy of the conformational ensemble (see Figure SI-21) as suggested by the study of Zapletal et al in 2020.^115^ However, it seems like it is still not enough to sufficiently modulate the phosphate/cation interaction, and that the ECC correction is a necessary addition to recover a more accurate conformational ensemble.

In order to assess the impact of the predictor choice, we also calculated SCSs using the PPM program^104^ (Tables SI-9 and SI-10 and Figure SI-22). PPM could not compute results for the *H*_*α*_ SCSs, but only *H*_*N*_. For all forcefields with the SCSs of *H*_*N*_, the observations made about the profiles are overall consistent between the PPM and SPARTA+ predictors, with only slight modifications at E3, A4, V16 and A17. The only discrepancy between the predictors is observed for the RMSE[H_*N*_], where PPM predicts A99-IPP closer to the experimental results while SPARTA+ designates A99. The predictor choice has thus a limited impact on the predictions and both predictors agree on an overall better agreement with experimental values of ECC-corrected forcefields over their full-charge parents.

## DISCUSSION

### ECC as a promising strategy to improve interactions between phosphorylated residues and cations. Impact on cation-mediated bridges

Comparing different popular force field combinations with experimental osmotic coefficient data on a small model system for doubly charged phosphate moieties, we show that mostly used non-polarizable force fields yield an unphysically strong interaction between cations and phosphorylated residues. This results in artefactual long-lived cation-mediated bridges between phosphorylated residues in MD simulations of phosphorylated peptides.

We showed that this overbinding artefact cannot be easily solved by a choice of water or ion force field, but is instead a common behavior for all non-polarizable force fields. In contrast, deriving ECC variants of the most popular force fields—where ionic charges are scaled by a factor 0.8 to account implicitly for electronic polarizability—efficiently eliminates these overbinding artefacts and yields excellent agreement with experimental osmotic coefficients, and a robustness across a broad concentration range (Figure 3).

When tested on the highly phosphorylated peptide 7PP, this results in much more extended conformations (Figure 5) and reduction of local curvature (Figure SI-18). These changes can be linked to a significant reduction of both the number and lifetime of cation-mediated bridges (*n*P-collabs) that are often reported in MD simulations of phosphorylated peptides, as we discussed in a previous work.^29^ The stability and occurence of such bridges seems to be strongly overestimated by standard force fields, with ECC variants all pointing to less common and much more transient formation of cation-mediated bridges between phosphorylated residues. The artefactually strong cation bridges may affect a lot of earlier simulation studies. For instance, de Bruyn et al. published very recently an exhaustive characterization of the conformational ensemble of *α*-synuclein and its phosphorylated form at S129.^116^ They reported that the forcefields DES-Amber with standard TIP4P-D^56^ and a99SB-disp with its modified TIP4P-D^117^ both display a phosphoserine that binds strongly to a sodium cation. This strong ion pair screens the electrostatic interactions of the phosphoresidue with the rest of the peptide, influencing the conformational ensemble of the phosphorylated form of *α*-synuclein. These examples, together with our results, draw a concerning picture not only for simulations of multiphosphorylated IDPs and peptides, but also for investigations of single phosphorylations with commonly employed full charge non-polarizable force fields. The overbinding of cations to phosphoresidues in classical forcefields could indeed prove problematic notably in the research of neurodegenerative diseases such as Alzheimer or Parkinson diseases for which abnormal phosphorylation patterns is a hallmark (Tau protein, *α*-synuclein respectively).

We show the potential of the ECC strategy to improve ion pairing behavior and allow for more realistic simulations of phosphorylated IDPs, without any increase in computational cost. While the ECC force field variants used here are certainly not as optimized and validated as common force fields, we note that the parameters we used for both ions and charged protein groups are very similar to those developed independently by Hui et al. and successfully employed for potassium channels simulations.^53^ While the development of ECC biomolecular force fields is still in its infancy, this lets us hope for a fast convergence of independent efforts.

A limitation of the present study resides in the difficulty to obtain a clear and quantitative experimental comparison to further confirm or tune the ECC parameters. Ideally, we would need to have experimental characterization of the conformational ensemble (e.g. radius of gyration) of short peptides with (one or several) or without phosphorylations, in different salt conditions (e.g. nature of the ions, concentration). Additional data on chemical shifts in systems with different conformational behavior would help us validate our models and pick the most accurate, provided we can train an accurate predictor of these chemical shifts.

### Limitations and shortcomings regarding the experimental comparison

This work underlines the challenges that still exist in the parameterization of non-bounded interactions in classical forcefields, and in doing so highlights several limitations and shortcomings for both experimental studies and computational modelling.

Computationally, the main hurdle remains the combination of forcefields used to represent protein dynamics. We assumed that the most significant bias induced in the peptide dynamics could be attributed solely to the phosphorylations and their interactions with ions, and that errors due to protein and water forcefields could be neglected. This is a strong assumption, since we observed in our previous study that even for unphosphorylated peptides, different forcefields do not necessarily converge towards the same behavior. ^29^ We had indeed noticed a stronger compaction from the C36 conformation ensemble compared to A99 for the R2 peptide in the unphosphorylated case. Testing of phosphorylation parameters will therefore require deeper investigation of the base protein forcefields as well.

Another issue when assessing local dynamics is that chemical shift predictors are not developed for phosphorylated amino-acids, which could degrade their predictive power. Although SPARTA+ and PPM predictions of SCSs (Figures 8, SI-13 and SI-14) tend to concur, it is hard to assess whether this results from an accurate prediction from both programs, or if both suffer from the same biases. Predictions were performed by specifying phosphoserines and phosphothreonines as regular serines and threonines. A recent study by Bakker et al. however suggests that the predictors struggle to make an accurate prediction for phosphoresidues.^118^ The overall discrepancy from the *H*_*N*_ SCSs is of the order of 0.4 ppm according to the RMSEs. This could be explained by an underestimation of the chemical shifts of phosphoresidues by the predictors, as the chemical shift of a phosphoserine and phosphothreonine are 9.13 ppm and 9.09 ppm respectively, compared to 8.39 ppm and 8.32 ppm for serine and threonine.^106,108^ Moreover, values closer to the experimental reference are values of unphosphorylated residues, while phosphoresidues display higher differences between the predicted and experimental value (Figure 8). Lastly on the computational side, simulations of disordered peptides and proteins are notoriously hard to converge, and the difficulty increases with peptide length. It is thus easier to obtain the conformational ensemble of shorter sequences, while experiments usually have a lower size limit in order to be able to accurately perform a measurement. This antagonistic relation to sequence length limits the possible comparisons between molecular dynamics simulations and experimental observables.

Experimentally, a first difficulty is the site-specific placement of phosphorylations. Experimentalists often need to employ phosphomimetic mutants by mutating the phosphosite to a glutamate residue, or to mutate unwanted phosphorylation sites to non-phosphorylatable residues in order for kinases to only target specific residues. ^119,120^ Confirming that these approaches offer a good approximation to the wild-type phosphorylated proteins can however prove challenging, and so becomes the comparison with MD simulations. ^121–123^ In addition, most experiments can only probe the average of observables at the global scale such as the hydrodynamic radius, which can be insensitive to local changes. To obtain information at the residue level in an IDP, the current method of choice is NMR spectroscopy. NMR experiments however present limitations which affect their ability to characterize cation/phosphorylations interactions. Low concentration or absence of salts is indeed preferred in order to improve the quality of the NMR acquisition. This explains why this study focuses on a single case with an extreme amount of phosphorylations, since it is this extreme composition which probably prompted the use of a high concentration of sodium phosphate as a buffer. Finally, the popularity of the sodium phosphate buffer in NMR experiments makes it difficult to assess the interaction between the cations and the phosphoresidues, since bulk phosphates could also be involved.

## CONCLUSION

We investigated the interaction between cations Na^+^ and K^+^ with phosphate groups for four classical forcefields through osmotic coefficient calculations, and all of them displayed concerning discrepancy with the experimental values. This led us to conclude that the interaction is too strong to be chemically realistic, and by extension that long-lasting cation-mediated bridges (*n*P-collabs) are most probably artefacts due to the the lack of electronic polarizability in the classical forcefields. We therefore tested the ECC charge-scaling correction scheme for phosphoresidues and bulk ions, also deriving ECC variants of the charged residues in C36 and A99 force fields. Simulations with these ECC variants show excellent agreement with experiment on the model phosphate compound, and were then applied to a 7-fold phosphorylated peptide from rhodopsin called 7PP. The ECC-corrected forcefields produce more extended conformations in average, and better reproduce experimental trends in secondary chemical shifts. We believe that the ECC strategy is very promising to solve the overstability of cation bridge and eliminate the associated artefacts—overcompaction of the conformational ensemble, increased stiffness—but these new protein forcefield should be carefully checked and validated —and if necessary adapted—before its usage can be routinely advised. An intermediate—though less consistent—possibility is to focus the ECC correction on bulk ions and phosphorylations, which should limit the possible artefacts. Among the classical, non rescaled force fields, the C36 forcefield with K^+^ bulk cations seems the less prone to overbinding artefacts, with the original parameters yielding a reasonable (though too low) osmotic coefficient compared to the experimental value. We hope through this study to convince both the computational and experimental communities that bulk ions play a major role in the conformational ensemble of phosphorylated peptides and IDPs, and that further work is needed to fully characterise the cation/phosphorylation interaction.

## Supporting information

Supplementary_informations_for_article_sticky_salts

## Data availability

Forcefield parameters, files and scripts required to perform osmotic coefficients, Δ*G* calculations, and molecular dynamics simulations of the 7PP peptide are deposited on Zenodo : https://doi.org/10.5281/zenodo.16980659

## Acknowledgement

The authors thank Dr. Josef Hritz and Dr. Vojtech Zapletal for sharing their TIP4P-D water model implementation for CHARMM36m; Dr. Kresten Lindorff-Larsen for his advice regarding the use of secondary chemical shifts, and Etienne Reboul for his help in developing analysis codes.

This work was supported by the ANR (MAGNETAU-ANR-21-CE29-0024) and the “Initiative d’Excellence” program from the French State (Grants “DYNAMO”, ANR-11-LABX-0011, and “CACSICE”, ANR-11-EQPX-0008). JPF acknowledges a PhD grant from the DYNAMO Labex. EDD acknowledges support from the ANR JCJC MUSIRICAT (ANR-22-CE29-0004-01). This project was provided with HPC computing and storage resources by GENCI (project A0170711021).

## Supporting Information Available

The following files are available free of charge.

- Supplementary_informations.pdf: contains the figures and tables listed as supplementary information in the paper, notably the partial charges and ionic parameters also available in the Zenodo repository : https://doi.org/10.5281/zenodo.16980659

## TOC Graphic

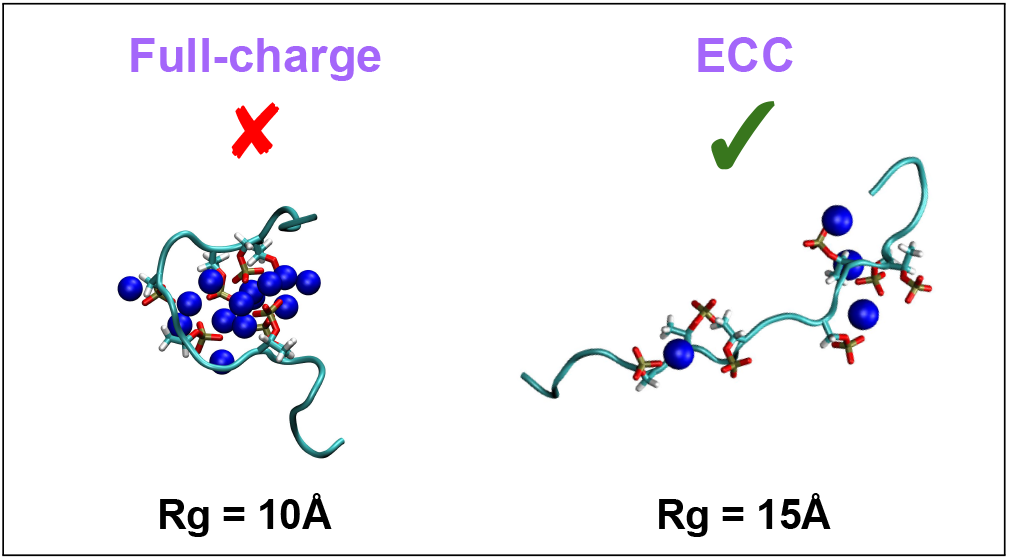

