## Supplementary_informations_for_article_sticky_salts for "Sticky salts: overbinding of monovalent cations to phosphorylations in all-atom forcefields"

### Supplementary Informations for Sticky salts: overbinding of monovalent cations to phosphorylations in all-atom forcefields

Jules Marien,[†,‡] Julie Puyo-Fourtine,[†,‡] Chantal Prévost,[†] Sophie Sacquin-Mora,[*,†] and Elise Duboué-Dijon[*,†]

[†] *Université Paris Cité, CNRS, Laboratoire de Biochimie Théorique, 13 rue Pierre et Marie Curie, 75005, Paris, France*

[‡] *Co-first authors*

Associated Zenodo repository : https://doi.org/10.5281/zenodo.16980659

| Ion | Forcefield | $\sigma$ (nm) | $\varepsilon$ (kJ/mol) | $q$ (e) |
|---|---|---|---|---|
| | Sengupta et al (A19) | 0.30326192 | 0.58382766 | 1.0 |
| | DES-Amber SF1.0 | 0.27830000 | 1.1694 | 1.0 |
| | Joung and Cheatham | 0.28330600 | 1.1692820 | 1.0 |
| **$K^+$** | **Joung and Cheatham - ECC** | **0.26914100** | **1.1692820** | **0.8** |
| | Beglov and Roux (C36, A18) | 0.314264522824 | 0.3640080 | 1.0 |
| | **Beglov and Roux - ECC (C36, A18)** | **0.298551290000** | **0.3640080** | **0.8** |
| | Beglov and Roux (A99) | 0.31426452 | 0.3640080 | 1.0 |
| | **Beglov and Roux - ECC (A99)** | **0.29855129** | **0.3640080** | **0.8** |
| | Sengupta et al (A19) | 0.26138968 | 0.12386075 | 1.0 |
| | DES-Amber SF1.0 | 0.20840000 | 0.7046 | 1.0 |
| | Joung and Cheatham | 0.21844800 | 0.7047430 | 1.0 |
| **$Na^+$** | **Joung and Cheatham - ECC** | **0.20752560** | **0.7047430** | **0.8** |
| | Beglov and Roux (C36, A18) | 0.251367073323 | 0.1962296 | 1.0 |
| | **Beglov and Roux - ECC (C36, A18)** | **0.230842990000** | **0.1962296** | **0.8** |
| | Beglov and Roux (A99) | 0.24299263 | 0.1962296 | 1.0 |
| | **Beglov and Roux - ECC (A99)** | **0.23084299** | **0.1962296** | **0.8** |
| | Sengupta et al (A19) | 0.42050419 | 2.8400519 | -1.0 |
| | DES-Amber SF1.0 | 0.47180000 | 0.0490 | -1.0 |
| | Joung and Cheatham | 0.49177600 | 0.04879170 | -1.0 |
| **$Cl^-$** | **Joung and Cheatham - ECC** | **0.46718700** | **0.04879170** | **-0.8** |
| | Beglov and Roux (C36, A18) | 0.404468018036 | 0.6276000 | -1.0 |
| | **Beglov and Roux - ECC (C36, A18)** | **0.384244619000** | **0.6276000** | **-0.8** |
| | Beglov and Roux (A99) | 0.40446802 | 0.6276 | -1.0 |
| | **Beglov and Roux - ECC (A99)** | **0.384244619** | **0.6276** | **-0.8** |

Table SI-1: Partial charges $q$, Lennard-Jones parameters $\sigma$ and $\epsilon$ of the ions considered in this study. Original parameters are in black, ECC-corrected parameters are highlighted in bold. Number of significant digits were kept as instructed in the forcefields.

| System | Contact (Å) | SShIP (Å) |
|---|---|---|
| C36, $Na^+$ | 0–4.4 | 4.4–7.1 |
| C36, $K^+$ | 0–4.5 | 4.5–7.0 |
| A99, $Na^+$ | 0–4.2 | 4.2–7.2 |
| A99, $K^+$ | 0–4.7 | 4.7–7.0 |

Table SI-2: Distance ranges used to define contact and solvent-shared ion pairs (SShIP) based on the first and second minima of the cation–phosphorus radial distribution functions.

| Forcefield combination | $Na^+$ | $Na^+$ ECC | $K^+$ | $K^+$ ECC |
|---|---|---|---|---|
| C36 + TIP3P* + BR ions | $0.11 \pm 0.02$ | | $0.68 \pm 0.02$ | |
| C36 + TIP4P-D + BR ions | $0.30 \pm 0.03$ | $0.78 \pm 0.03$ | $0.52 \pm 0.02$ | $0.81 \pm 0.03$ |
| A18 + TIP3P-FB + JC ions | $0.11 \pm 0.02$ | | $0.38 \pm 0.03$ | |
| A19 + OPC + SM ions | $0.59 \pm 0.03$ | | $0.65 \pm 0.03$ | |
| A99 + TIP4P-D + BR ions | $0.38 \pm 0.03$ | $0.80 \pm 0.02$ | $0.58 \pm 0.02$ | $0.83 \pm 0.02$ |
| A99 + TIP4P-D + DES ions | $0.27 \pm 0.03$ | | $0.51 \pm 0.02$ | |
| A99 + TIP4P-D + JC ions | $0.23 \pm 0.03$ | $0.78 \pm 0.04$ | $0.52 \pm 0.03$ | $0.78 \pm 0.02$ |

Table SI-3: Osmotic coefficients $\phi$ calculated for the different combinations of forcefields for $2Na^+HPO_4^{2-}$ and $2K^+HPO_4^{2-}$ at 0.2M concentration.

| Forcefield combination | 0.2M | 0.5M | 0.8M |
|---|---|---|---|
| C36 | $0.11 \pm 0.02$ | $0.04 \pm 0.01$ | $0.05 \pm 0.01$ |
| C36 ECC | $0.78 \pm 0.03$ | $0.69 \pm 0.01$ | $0.64 \pm 0.02$ |
| A99 | $0.38 \pm 0.03$ | $0.18 \pm 0.02$ | $0.12 \pm 0.02$ |
| A99 ECC | $0.78 \pm 0.04$ | $0.70 \pm 0.02$ | $0.66 \pm 0.02$ |

Table SI-4: Osmotic coefficients $\phi$ calculated for the different combinations of forcefields for $2Na^+HPO_4^{2-}$ at different concentrations.

| Forcefield combination | 0.2M | 0.5M | 0.8M |
|---|---|---|---|
| C36 | $0.68 \pm 0.02$ | $0.48 \pm 0.02$ | $0.34 \pm 0.02$ |
| C36 ECC | $0.81 \pm 0.03$ | $0.77 \pm 0.02$ | $0.75 \pm 0.02$ |
| A99 | $0.58 \pm 0.02$ | $0.48 \pm 0.02$ | $0.42 \pm 0.02$ |
| A99 ECC | $0.78 \pm 0.02$ | $0.71 \pm 0.02$ | $0.70 \pm 0.02$ |

Table SI-5: Osmotic coefficients $\phi$ calculated for the different combinations of forcefields for $2K^+HPO_4^{2-}$ at different concentrations.

| **Forcefield** | **Total** | | | **contact ion pair** | | | **solvent-shared** | | |
|---|---|---|---|---|---|---|---|---|---|
| | rep1 | rep2 | rep3 | rep1 | rep2 | rep3 | rep1 | rep2 | rep3 |
| A99 | −5.6 | −5.7 | −5.9 | −4.3 | −4.5 | −4.8 | −3.3 | −3.3 | −3.5 |
| A99 ECC | −4.0 | −3.8 | −4.2 | −2.3 | −2.1 | −2.4 | −2.3 | −2.1 | −2.5 |
| C36 | −4.0 | −3.9 | −3.6 | −2.8 | −2.7 | −2.1 | −1.6 | −1.6 | −1.5 |
| C36 ECC | −2.1 | −2.0 | −2.1 | 0.82 | 0.83 | 0.77 | −1.2 | −1.0 | −1.2 |

Table SI-6: Binding free energies (kJ/mol) between $HPO_4^{2-}$ and $K^+$ obtained for each replica of the calculation, in each binding mode and with all tested force field combinations.

| **Forcefield** | **Total** | | | **contact ion pair** | | | **solvent-shared** | | |
|---|---|---|---|---|---|---|---|---|---|
| | rep1 | rep2 | rep3 | rep1 | rep2 | rep3 | rep1 | rep2 | rep3 |
| A99 | −7.1 | −6.6 | −6.5 | −5.5 | −5.4 | −5.0 | −5.2 | −4.3 | −4.5 |
| A99 ECC | −3.8 | −3.9 | −3.4 | −1.0 | −0.9 | −0.5 | −2.8 | −3.0 | −2.5 |
| C36 | −5.8 | −5.7 | −5.9 | −5.2 | −5.2 | −5.3 | −2.0 | −1.5 | −1.9 |
| C36 ECC | −2.3 | −2.5 | −2.4 | 0.5 | 0.4 | 0.3 | −1.3 | −1.5 | −1.4 |

Table SI-7: Binding free energies (kJ/mol) between $HPO_4^{2-}$ and $Na^+$ obtained for each replica of the calculation, in each binding mode and with all tested force field combinations.

| **$\Delta G$ contribution (kJ/mol)** | **replicates** | | |
|---|---|---|---|
| $\Delta G^{*}_{bulk}$ | 280.7 | 280.9 | 280.6 |
| $\Delta G_{\mathrm{PBC}}$ | −1.4 | −1.4 | −1.4 |
| $\Delta G^{\mathrm{V}^{\circ}\to\mathrm{rest}}_{\mathrm{decoupled}}$ | 3.2 | 3.2 | 3.2 |
| $\Delta G^{*}_{site}$ | 286.9 | 287.3 | 287.2 |
| $\Delta G^{\circ}_{\mathrm{bind}}$ | −4.3 | −4.5 | −4.8 |

Table SI-8: Detailed free energy contributions for the calculation of binding free energies between $HPO_4^{2-}$ and $K^+$ for the contact ion pair and with the A99 force field, selected as an exemple.

| **Forcefields** | **RMSE[$\mathbf{H}_N$] (ppm)** | **$r$[$\mathbf{H}_N$] (ppm)** |
|---|---|---|
| C36 | 0.53 | 0.78 |
| C36+TIP4P-D | 0.39 | 0.83 |
| C36-IP | **0.36** | **0.87** |
| C36-IPP | 0.39 | 0.84 |

Table SI-9: RMSE and Pearson's correlation coefficients $r$ between the forcefield predictions of C36, C36+TIP4P-D, C36-IP and C36-IPP, and the experimental values for the SCS of $H_N$ of peptide 7PP with the PPM predictor. Best values are highlighted in bold.

| **Forcefields** | **RMSE[$\mathbf{H}_N$] (ppm)** | $r$[$\mathbf{H}_N$] **(ppm)** |
|---|---|---|
| A99 | 0.46 | 0.82 |
| A99-IP | 0.47 | **0.91** |
| A99-IPP | **0.44** | 0.88 |

Table SI-10: RMSE and Pearson's correlation coefficients $r$ between the forcefield predictions of A99, A99-IP and A99-IPP, and the experimental values for the SCS of $H_N$ of peptide 7PP with the PPM predictor. Best values are highlighted in bold.

Table SI-11: Average over all residues of the Local Curvatures (LCs) for the 7PP peptide simulations.

| | **C36** | **C36 ECC-IP** | **C36 ECC-IPP** | **A99** | **A99 ECC-IP** | **A99 ECC-IPP** |
|---|---|---|---|---|---|---|
| **Replica 1** | 0.14 | 0.11 | 0.11 | 0.16 | 0.15 | 0.15 |
| **Replica 2** | 0.15 | 0.11 | 0.10 | 0.13 | 0.15 | 0.15 |
| **Replica 3** | 0.15 | 0.11 | 0.12 | 0.12 | 0.14 | 0.14 |

Table SI-12: Average over all residues of the Local Fluctuations (LFs) for the 7PP peptide simulations.

| | **C36** | **C36 ECC-IP** | **C36 ECC-IPP** | **A99** | **A99 ECC-IP** | **A99 ECC-IPP** |
|---|---|---|---|---|---|---|
| **Replica 1** | 0.04 | 0.04 | 0.05 | 0.05 | 0.06 | 0.06 |
| **Replica 2** | 0.04 | 0.04 | 0.04 | 0.06 | 0.06 | 0.06 |
| **Replica 3** | 0.04 | 0.04 | 0.05 | 0.05 | 0.06 | 0.06 |

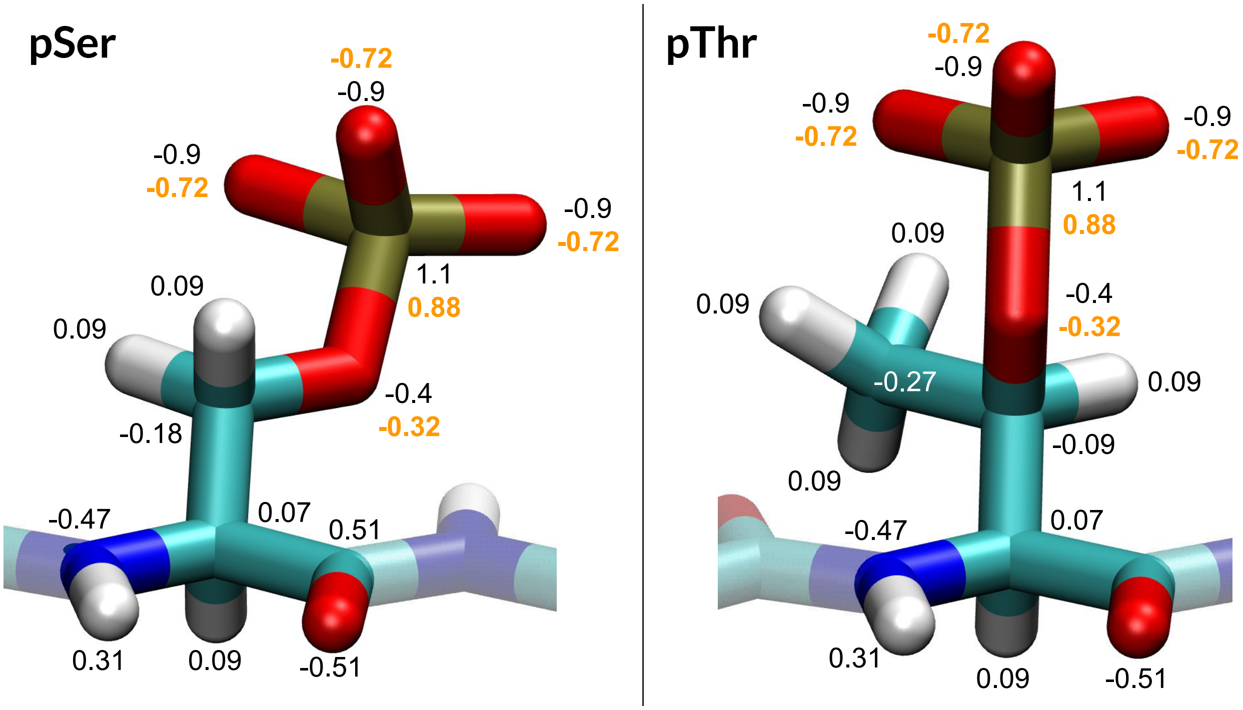


Figure SI-1: Partial charges of forcefield C36 for phosphoserine and phosphothreonine. Original parameters are in black or white, ECC-corrected parameters are in orange.

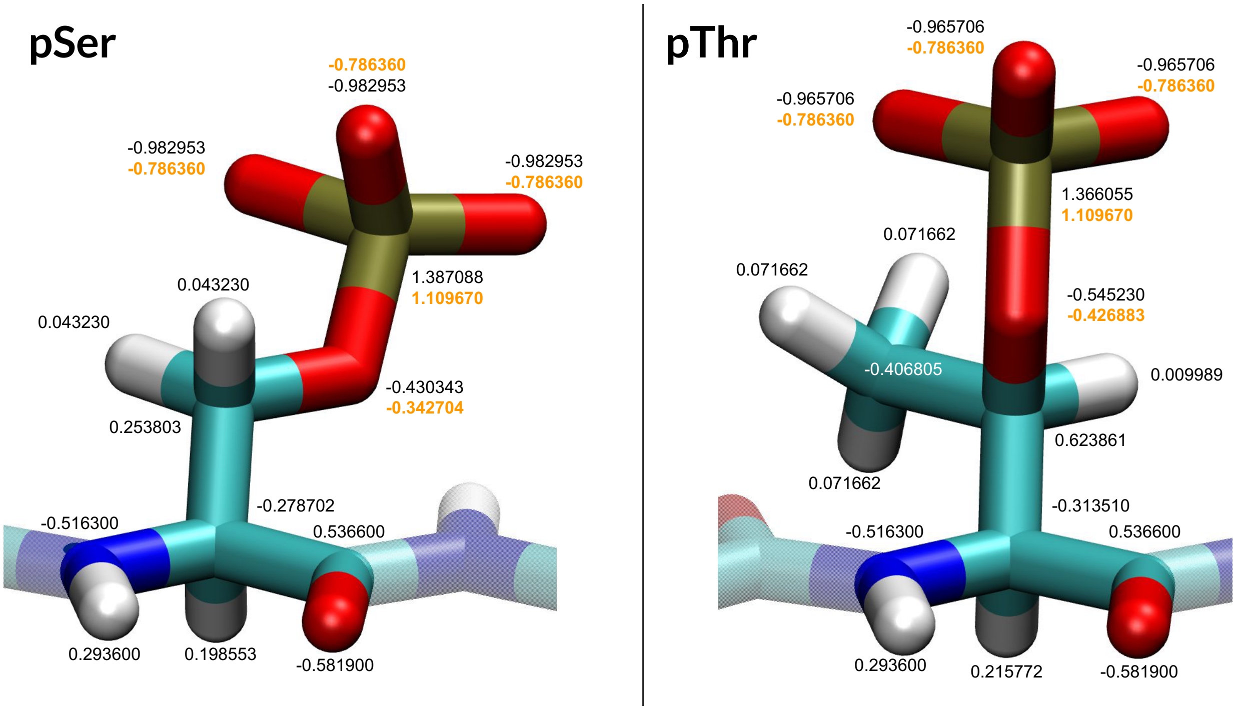


Figure SI-2: Partial charges of forcefield A99 for phosphoserine and phosphothreonine. Original parameters are in black or white, ECC-corrected parameters are in orange.

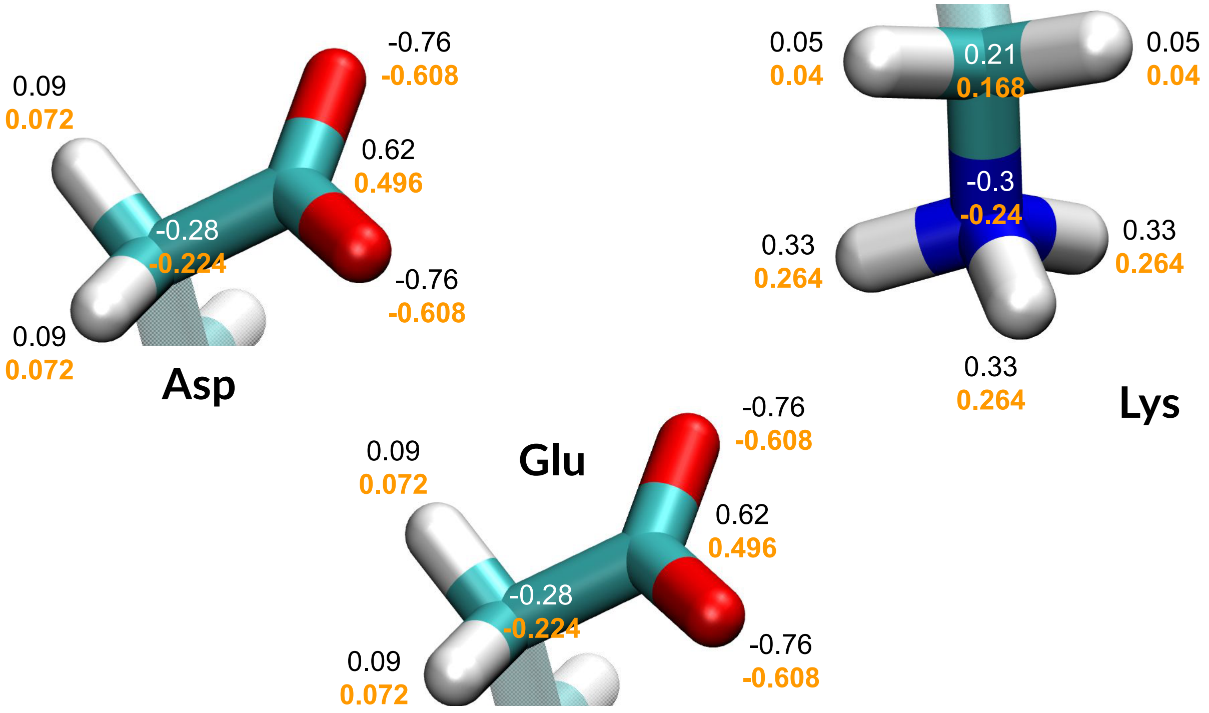


Figure SI-3: Partial charges of forcefield C36 for moieties carrying the charge of residues Asp, Glu and Lys. Original parameters are in black or white, ECC-corrected parameters are in orange.

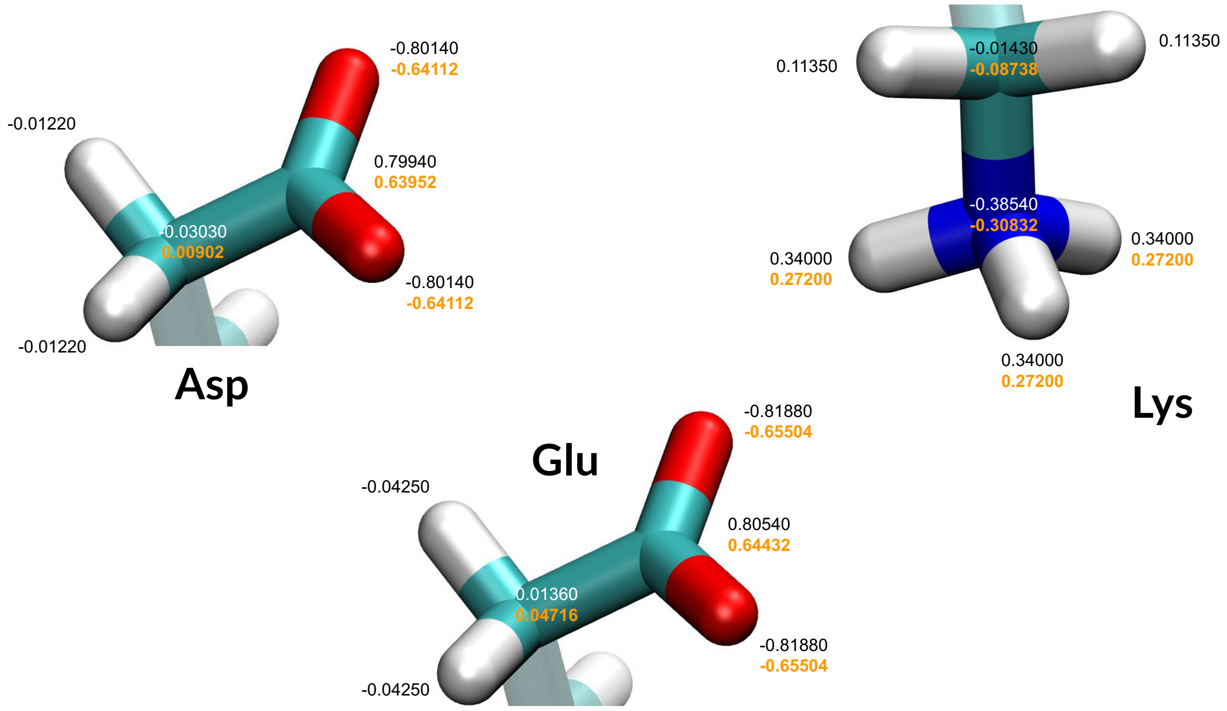


Figure SI-4: Partial charges of forcefield A99 for moieties carrying the charge of residues Asp, Glu and Lys. Original parameters are in black or white, ECC-corrected parameters are in orange.

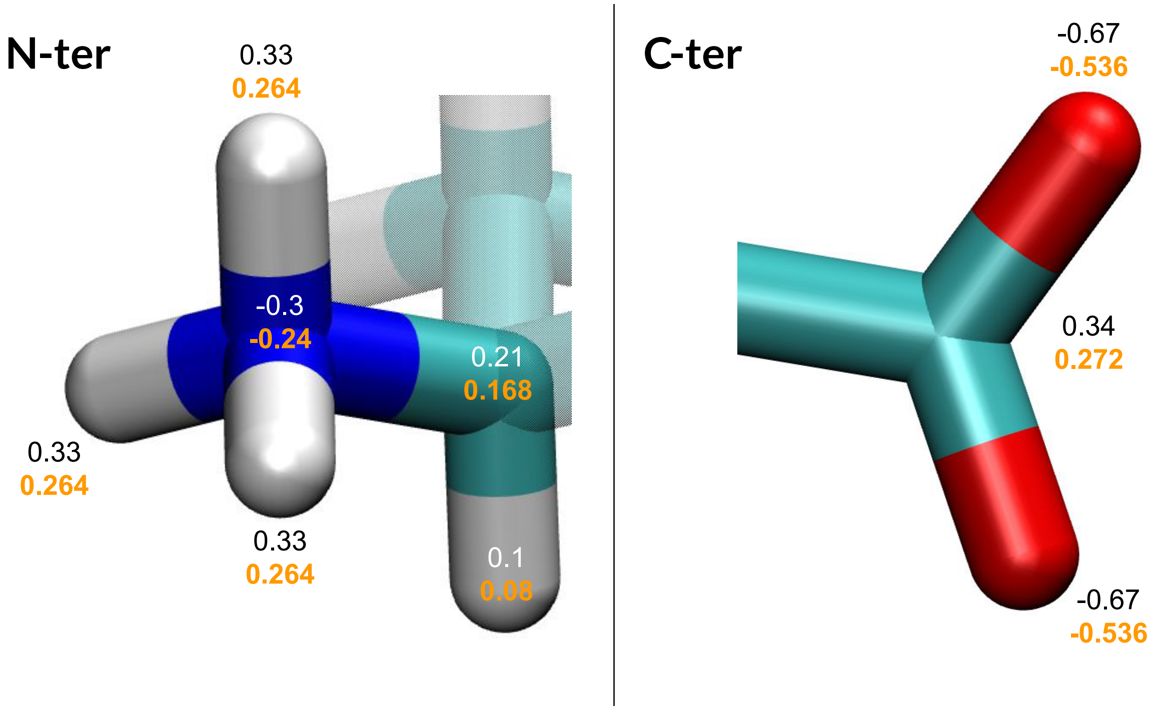


Figure SI-5: Partial charges of forcefield C36 for N-terminal and C-terminal moieties. Original parameters are in black or white, ECC-corrected parameters are in orange.

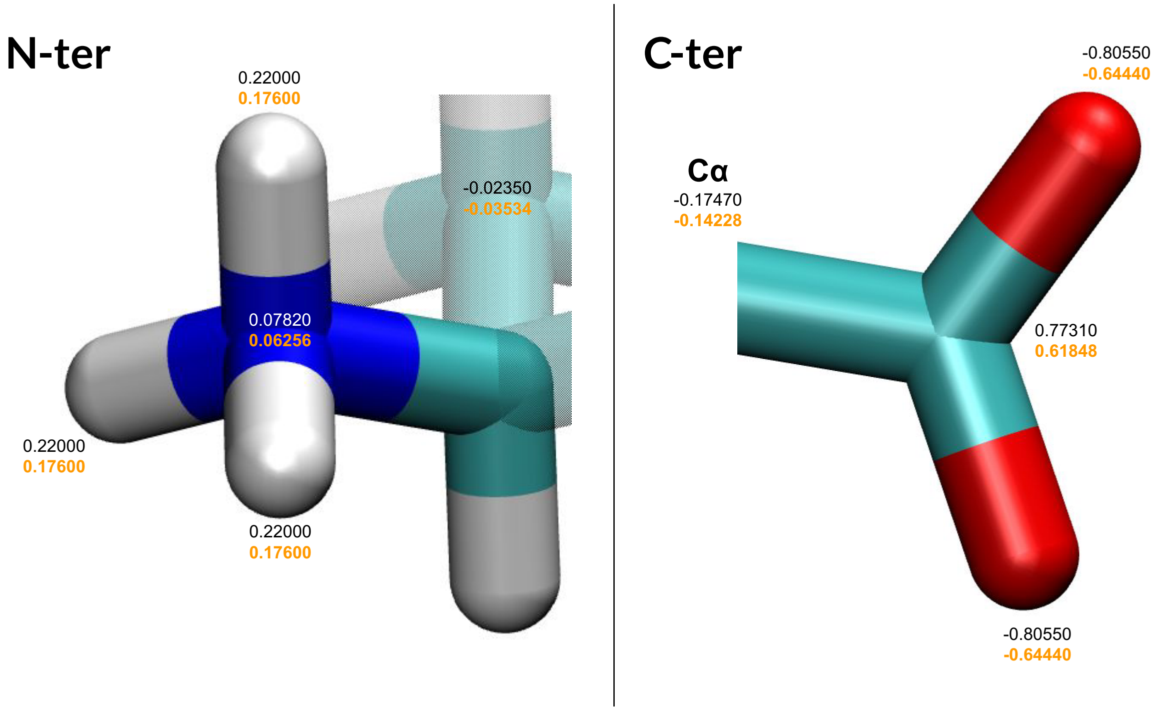


Figure SI-6: Partial charges of forcefield A99 for N-terminal and C-terminal moieties. Original parameters are in black or white, ECC-corrected parameters are in orange.

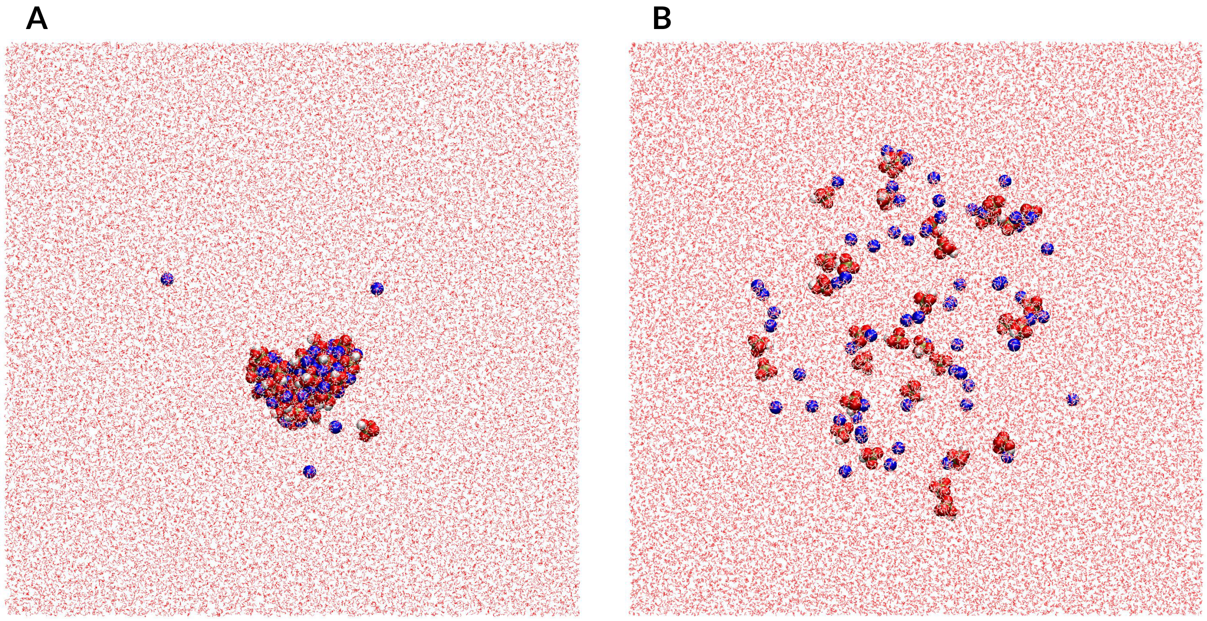


Figure SI-7: Last frame of the first replica of simulations at 0.2mol.L$^{-1}$ of $2Na^+HPO_4^{2-}$ for C36 (A) and ECC-C36 (B).

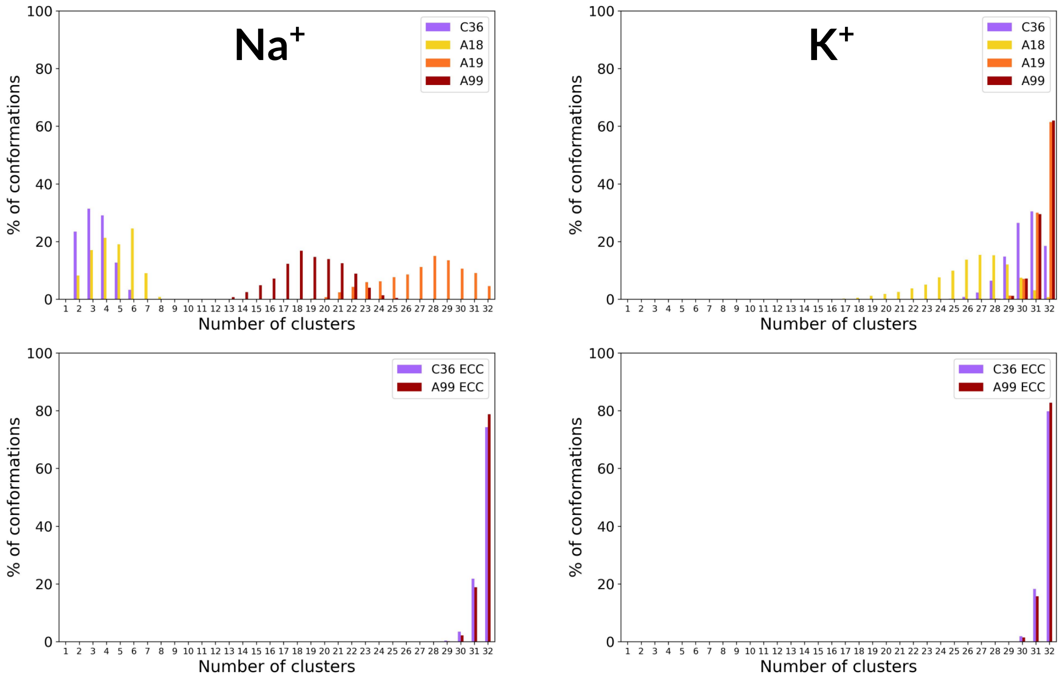


Figure SI-8: Number of clusters formed by the 32 $HPO_4^{2-}$ molecules when interacting with $Na^+$ (left) or $K^+$ cations during the simulations for the calculations of osmotic coefficients. 1 cluster means that all $HPO_4^{2-}$ molecules aggregated, 32 that none aggregated. Cluster cutoff was set at 0.6nm and all 10 replicas were cumulated.

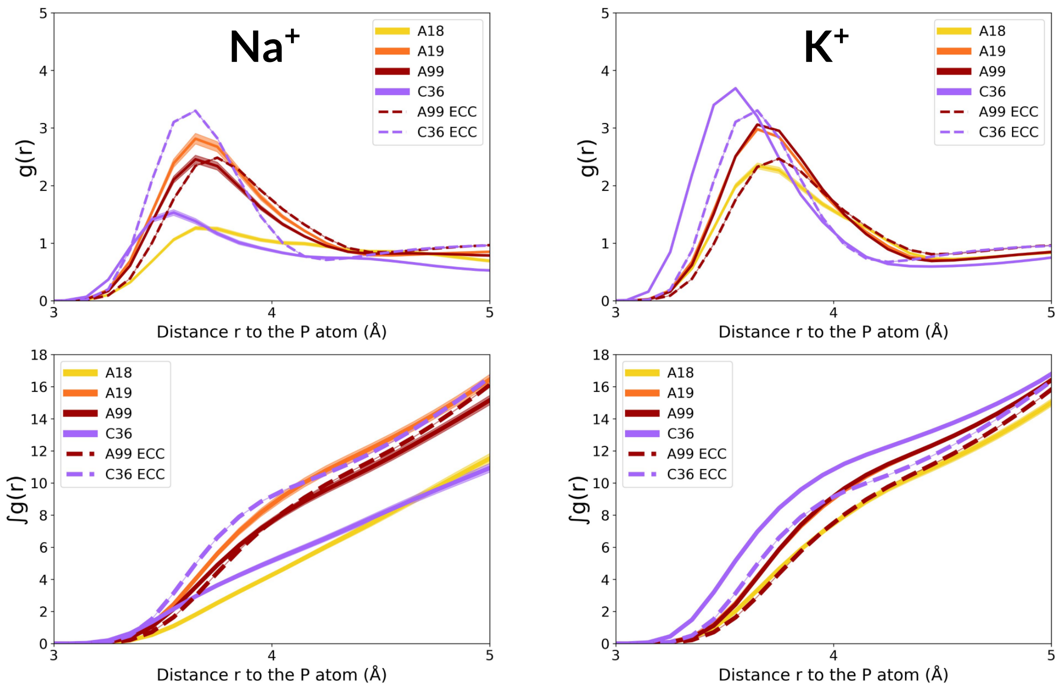


Figure SI-9: Radial distribution functions g(r) (upper panels) and their integrals over distance (lower panels) of the water oxygens around the phosphorus atoms contained in the 32 $HPO_4^{2-}$ molecules when interacting with $Na^+$ (left) or $K^+$ cations during the simulations for the calculations of osmotic coefficients. ECC-corrected forcefields are represented with dashed lines. The main line of each figure represents the average over the 10 replicas, and represented as a transparent background with proportional thickness are the standard deviations at each bin (increment of 0.1 Å).

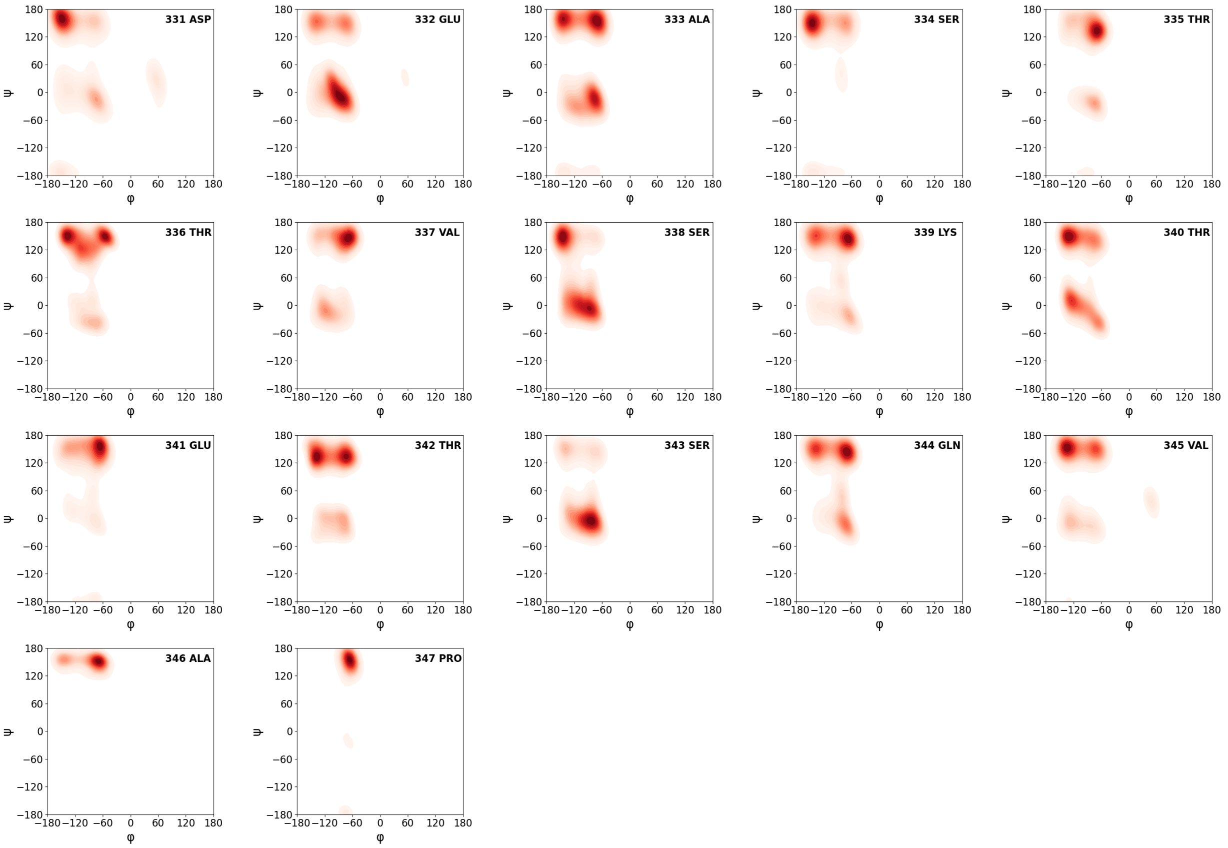


Figure SI-10: Ramachandran plots for the residues of 7PP simulated with the A99 forcefield. The plots combine all three replicas.

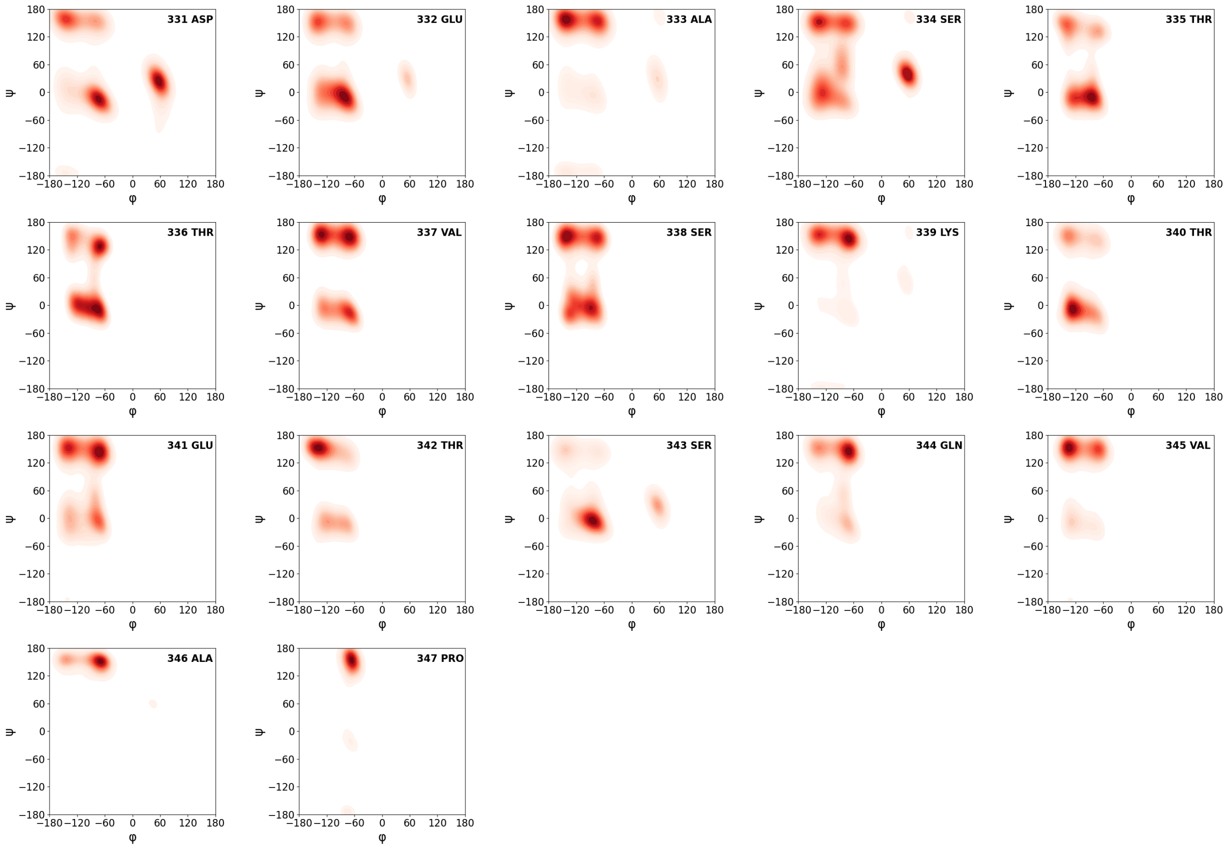


Figure SI-11: Ramachandran plots for the residues of 7PP simulated with the A99 ECC-IP forcefield. The plots combine all three replicas.

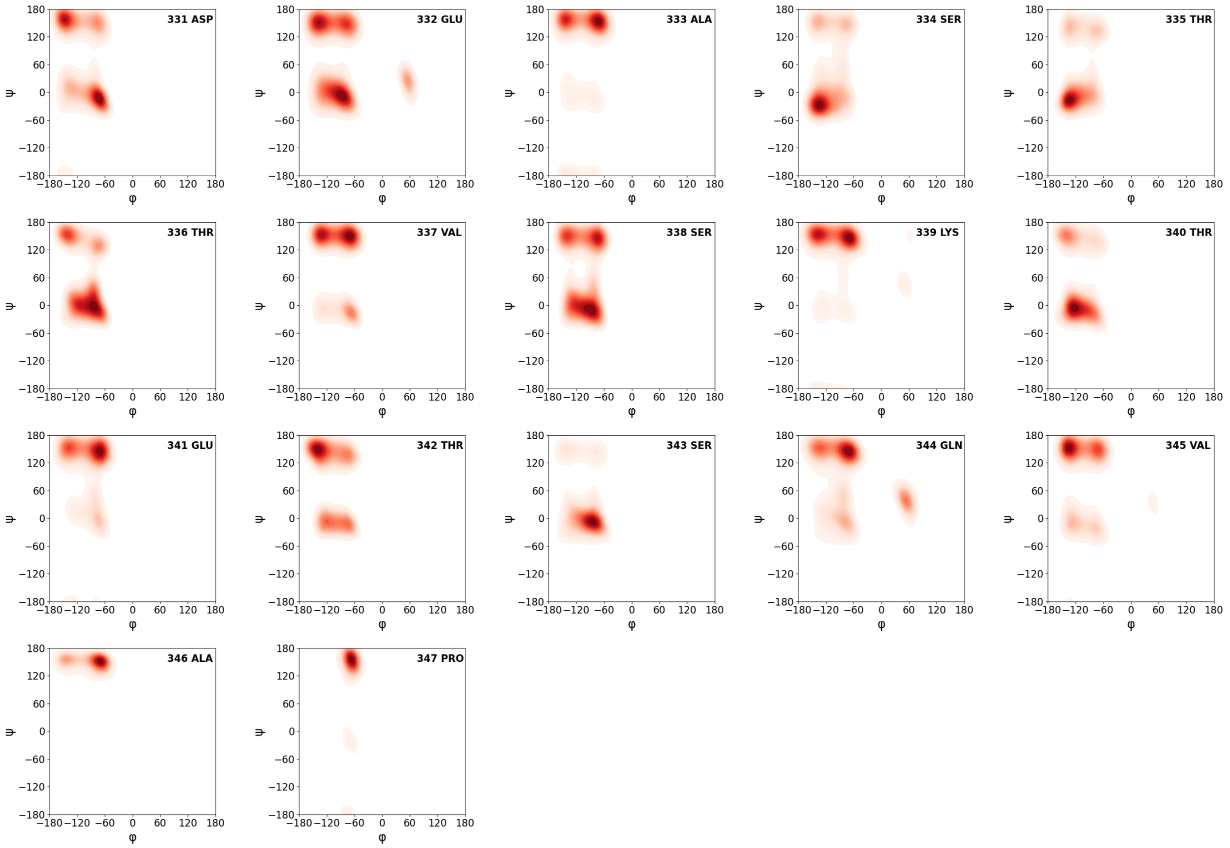


Figure SI-12: Ramachandran plots for the residues of 7PP simulated with the A99 ECC-IPP forcefield. The plots combine all three replicas.

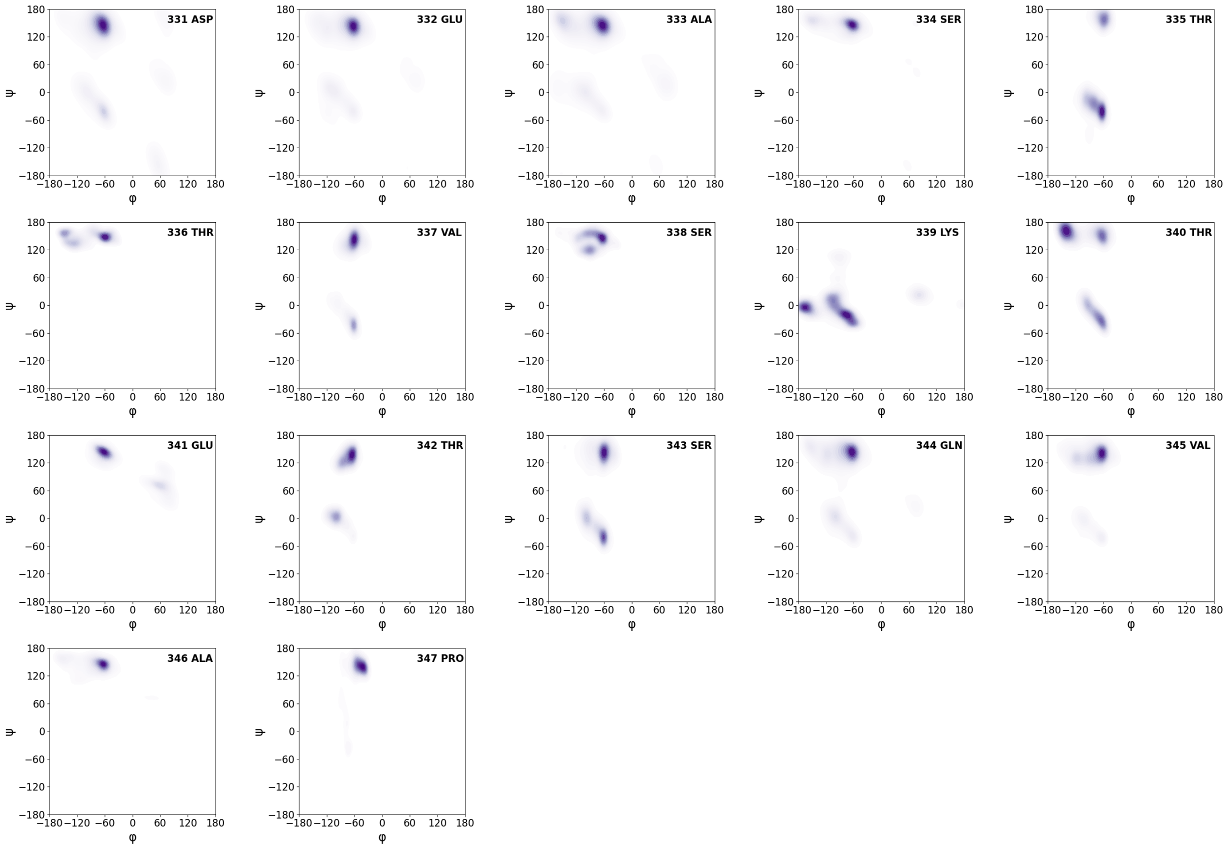


Figure SI-13: Ramachandran plots for the residues of 7PP simulated with the C36 forcefield. The plots combine all three replicas.

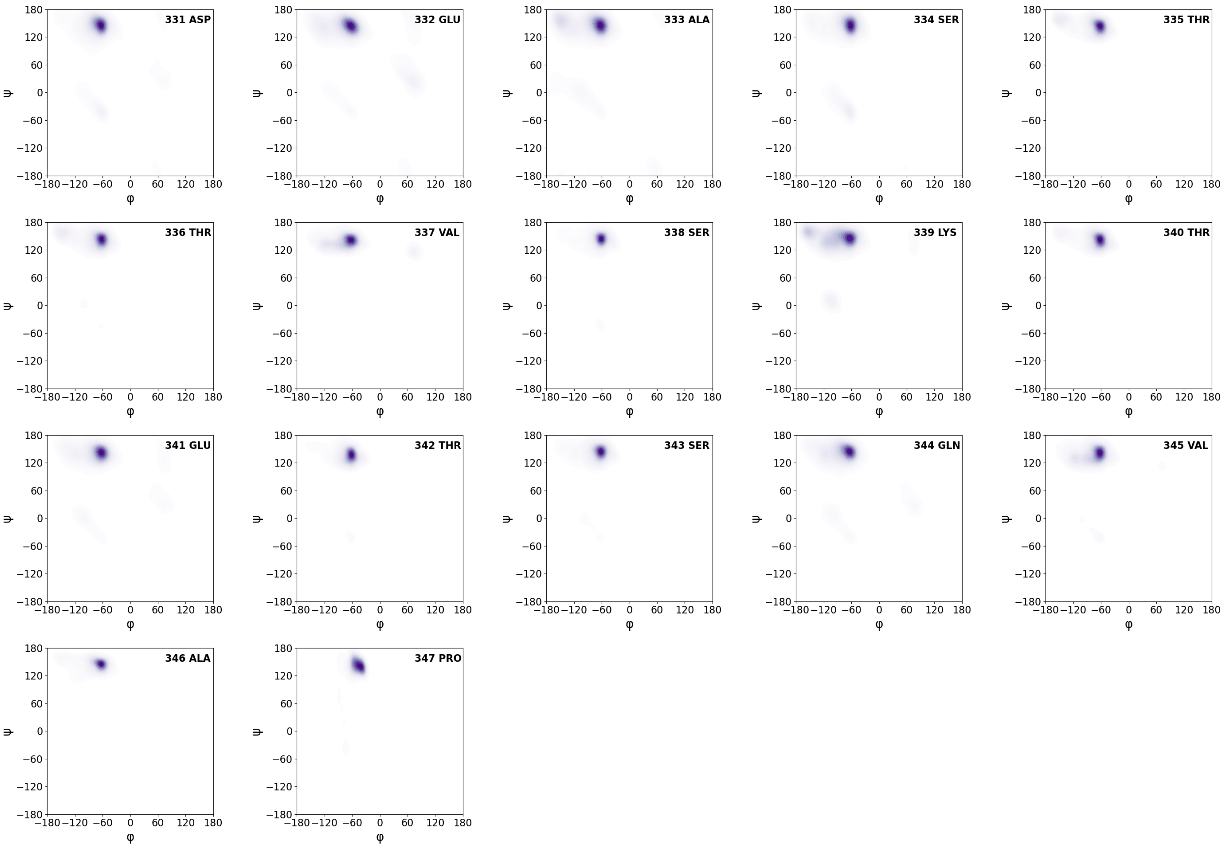


Figure SI-14: Ramachandran plots for the residues of 7PP simulated with the C36 ECC-IP forcefield. The plots combine all three replicas.

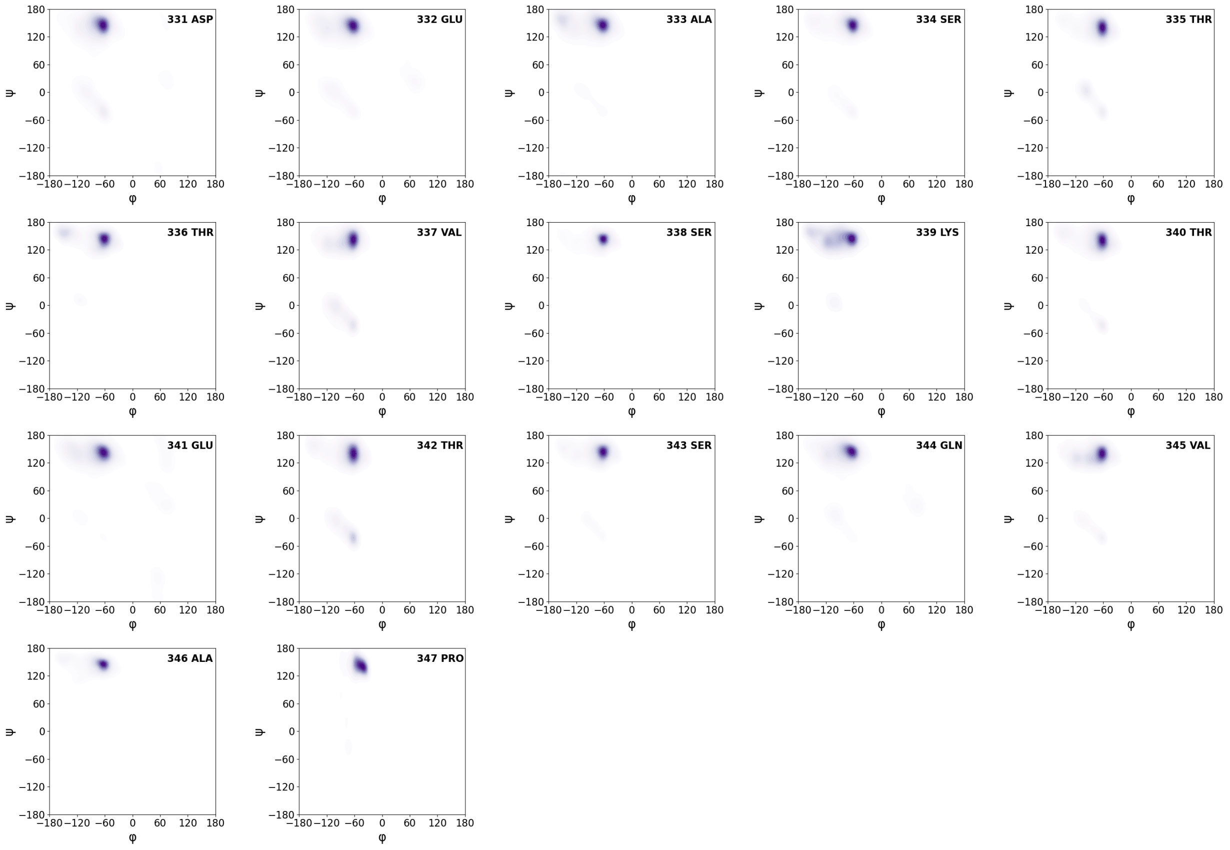


Figure SI-15: Ramachandran plots for the residues of 7PP simulated with the C36 ECC-IPP forcefield. The plots combine all three replicas.

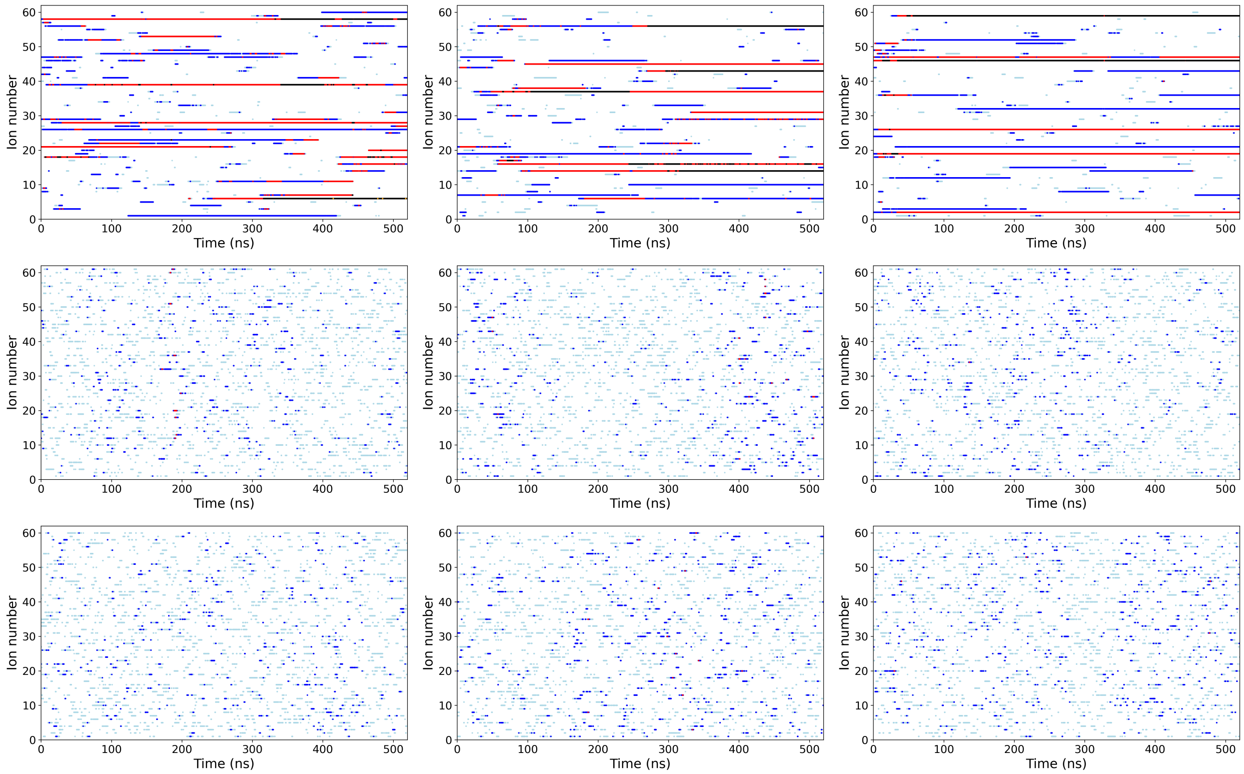


Figure SI-16: Presence of $Na^+$ as a function of time near the phosphates of the phosphorylations of the 7PP peptide for the C36 forcefield. If the cation is near one phosphate ($\leq$ 4Å to the phosphorus atom), a light blue dot is displayed. 2P-collabs are in dark blue, 3P-collabs in red, 4P-collabs in black and 5P-collabs and higher in orange. The first row corresponds to the three replicas of the C36 forcefield, the second to the replicas of the C36 ECC-IP forcefield and the third to the replicas of the C36 ECC-IPP forcefield.

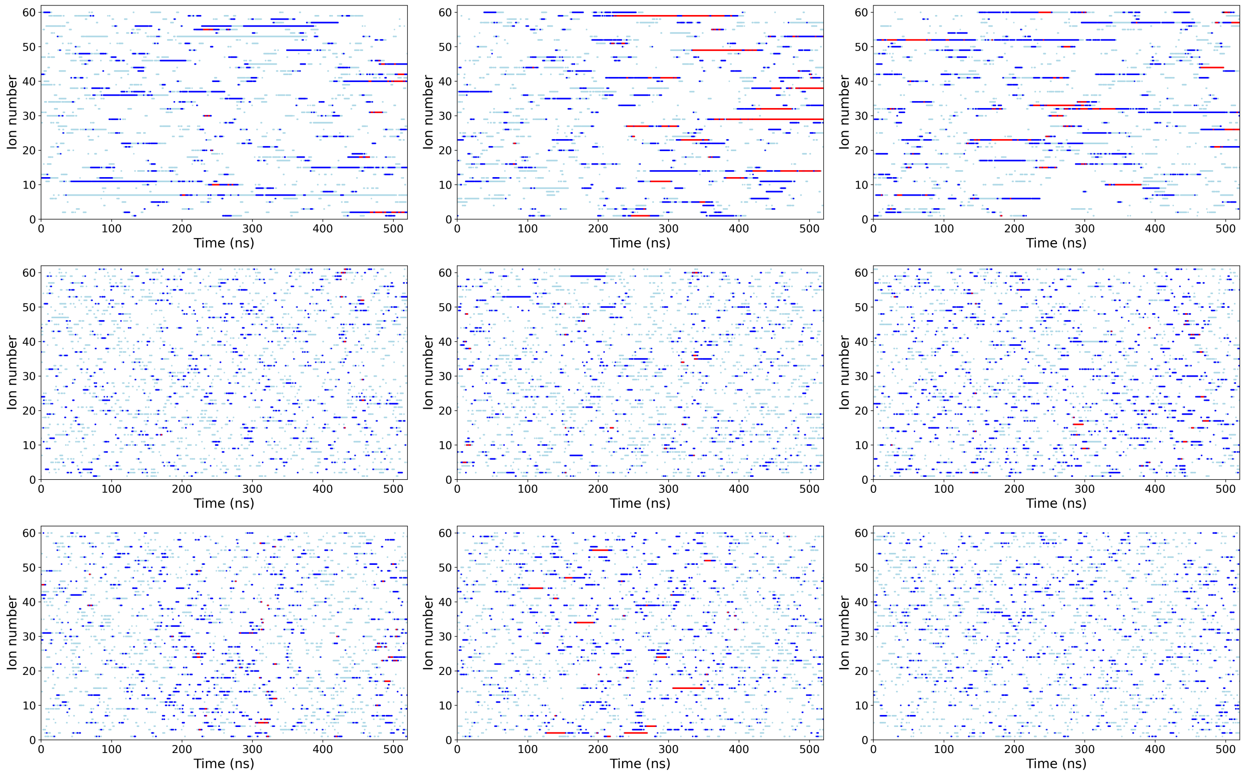


Figure SI-17: Presence of $Na^+$ as a function of time near the phosphates of the phosphorylations of the 7PP peptide for the A99 forcefield. If the cation is near one phosphate ($\leq 4$Å to the phosphorus atom), a light blue dot is displayed. 2P-collabs are in dark blue, 3P-collabs in red, 4P-collabs in black and 5P-collabs and higher in orange. The first row corresponds to the three replicas of the A99 forcefield, the second to the replicas of the A99 ECC-IP forcefield and the third to the replicas of the A99 ECC-IPP forcefield.

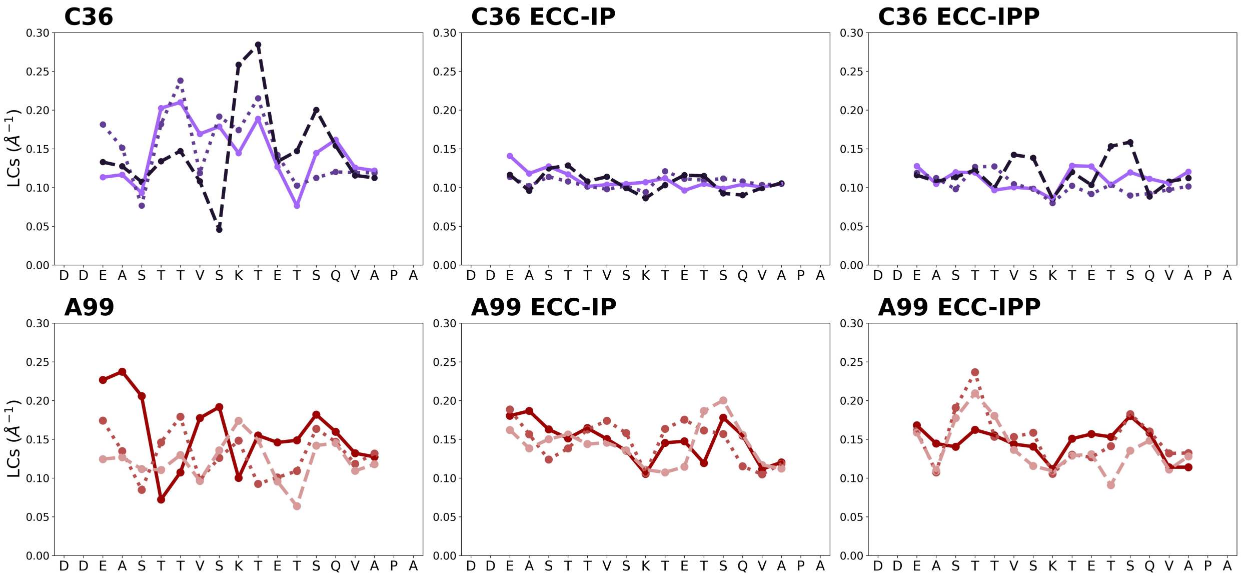


Figure SI-18: Local Curvatures (LCs) per replica of the 7PP peptide depending on the forcefields (C36 on the top row, A99 on the bottom). The first replica is in full line, the second replica in dashed line and the third replica in dotted line.

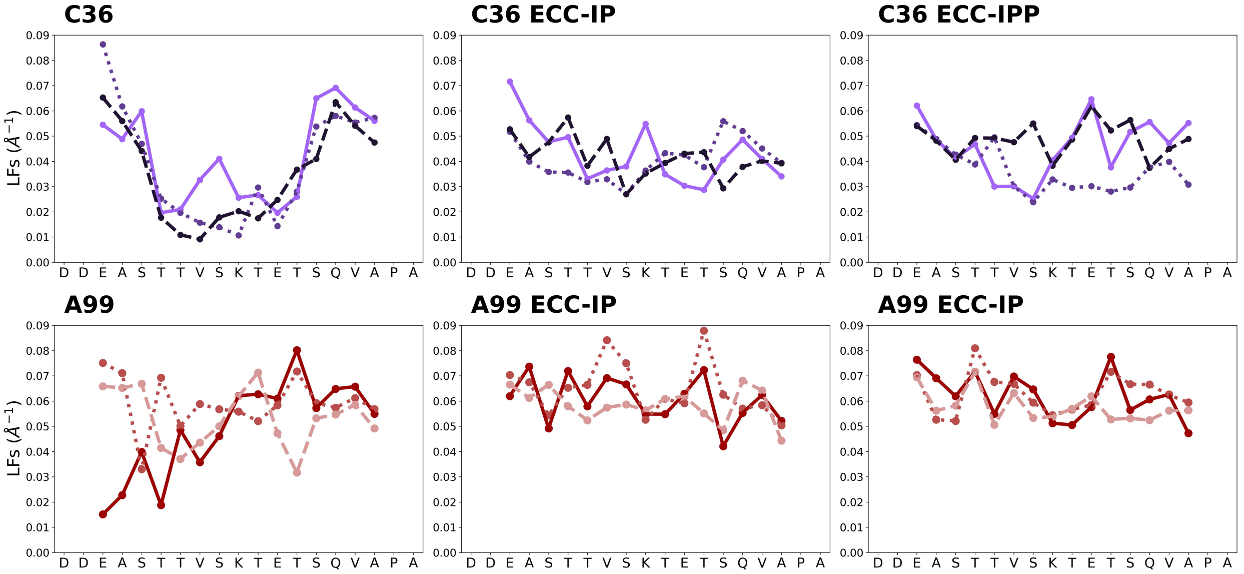


Figure SI-19: Local Flexibilities (LFs) per replica of the 7PP peptide depending on the forcefields (C36 on the top row, A99 on the bottom). The first replica is in full line, the second replica in dashed line and the third replica in dotted line.

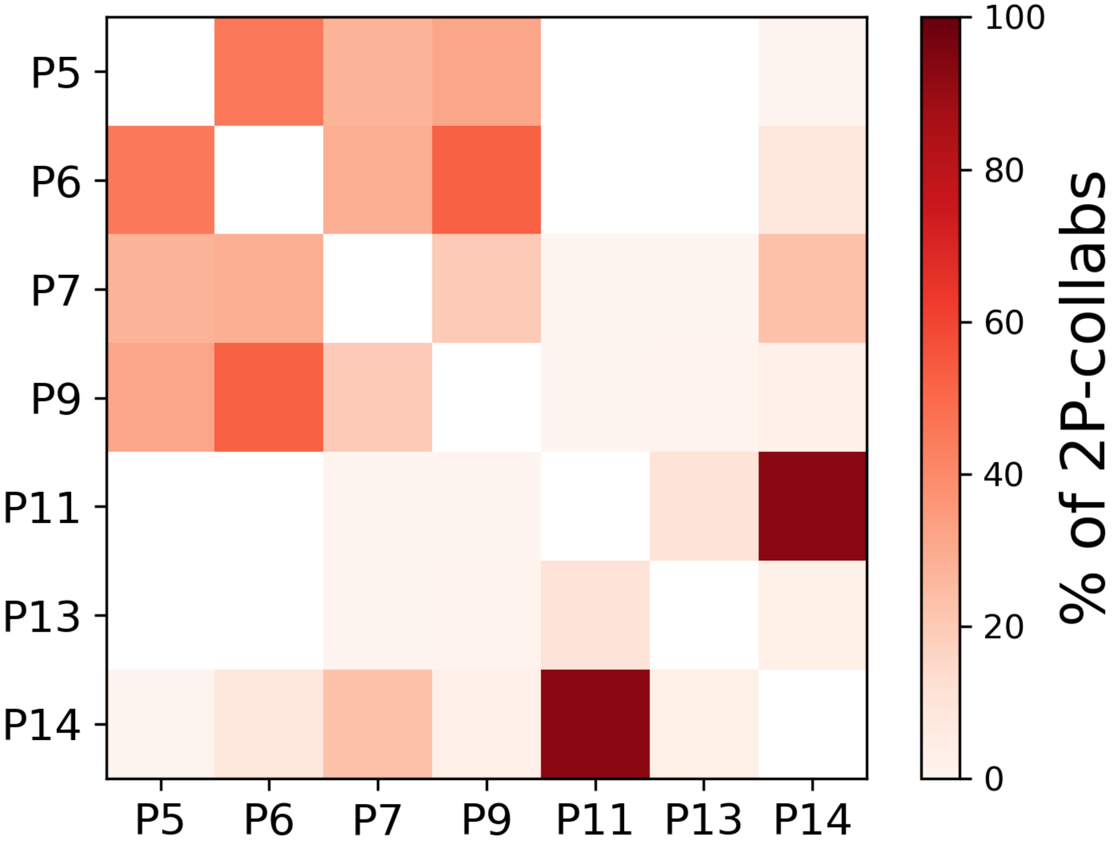


Figure SI-20: 2P-collab counts between the phosphoresidues of the C36 + TIP4P-D simulations of 7PP.

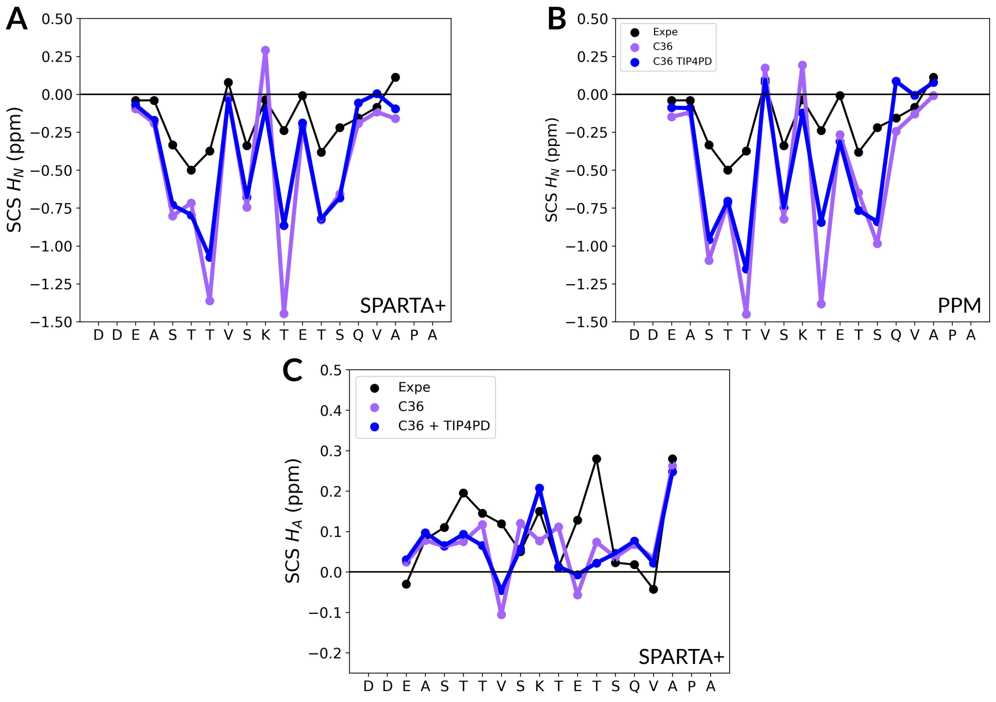


Figure SI-21: Secondary Chemical Shifts (SCS) of $H_N$ from the experimental data (in black) and from computation of the original C36 (in deep purple) against C36 with the TIP4P-D water model (in blue). A) $H_N$ predictions with SPARTA+, B) $H_N$ predictions with PPM, C) $H_A$ predictions with SPARTA+

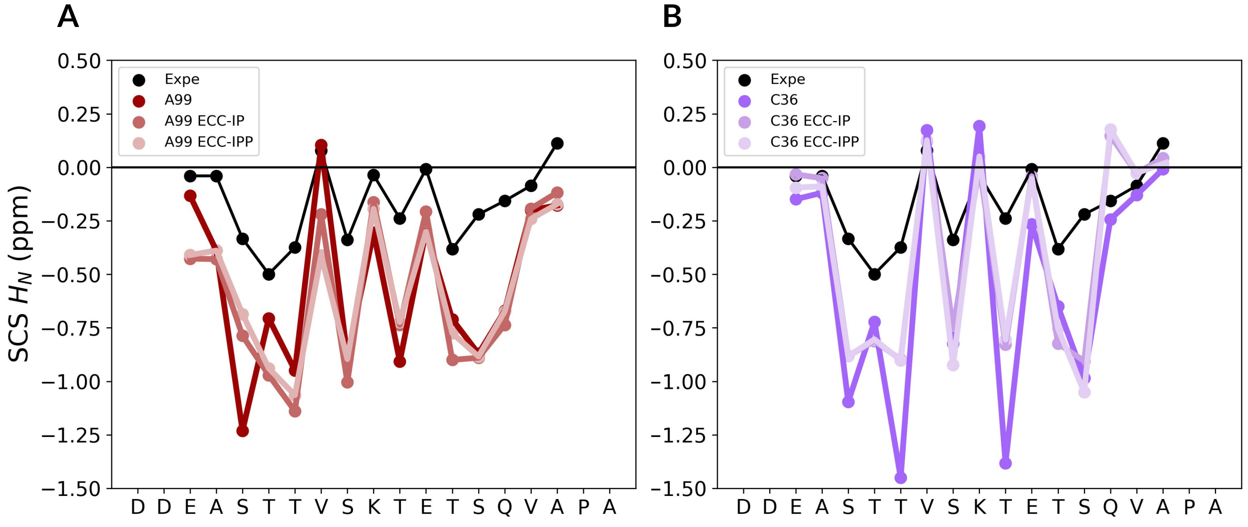


Figure SI-22: Secondary Chemical Shifts (SCS) of $H_N$ from the experimental data (in black) and from computation using PPM. A) A99, B) C36
